# MINTIE: identifying novel structural and splice variants in transcriptomes using RNA-seq data

**DOI:** 10.1101/2020.06.03.131532

**Authors:** Marek Cmero, Breon Schmidt, Ian J. Majewski, Paul G. Ekert, Alicia Oshlack, Nadia M. Davidson

## Abstract

Genomic rearrangements can modify gene function by altering transcript sequences, and have been shown to be drivers in both cancer and rare diseases. Although there are now many methods to detect structural variants from Whole Genome Sequencing (WGS), RNA sequencing (RNA-seq) remains under-utilised as a technology for the detection of gene altering structural variants. Calling fusion genes from RNA-seq data is well established, but other transcriptional variants such as fusions with novel sequence, tandem duplications, large insertions and deletions, and novel splicing are difficult to detect using existing approaches.

To identify all types of variants in transcriptomes, we developed MINTIE, an integrated pipeline for RNA-seq data. We take a reference free approach, which combines de novo assembly of transcripts with differential expression analysis, to identify up-regulated novel variants in a case sample.

We validated MINTIE on simulated and real data sets and compared it with eight other approaches for finding novel transcriptional variants. We found MINTIE was able to detect >85% of variants while no other method was able to achieve this.

We applied MINTIE to RNA-seq data from a cohort of acute lymphoblastic leukemia (ALL) patient samples and identified several clinically relevant variants, including a recurrent unpartnered fusion involving the tumour suppressor gene RB1, and variants in ALL-associated genes: tandem duplications in IKZF1 and PAX5, and novel splicing in ETV6. We further demonstrate the utility of MINTIE to identify rare disease variants using RNA-seq, including the discovery of an inter-chromosomal translocation in the DMD gene in a patient with muscular dystrophy. We posit that MINTIE will be able to identify new disease variants across a range of cancers and other disease types.

## Introduction

Rearrangements of the genome can disrupt or modify gene function and have been implicated as the causal event in disease. In cancer, somatic genomic rearrangements are common and can alter the genomic landscape to drive oncogenesis and cancer progression^1–3^. Whole genome sequencing (WGS) has successfully been used to detect structural variants (SVs) and profile their frequency in cancers^4^ and rare diseases^5,6^. However, clinically relevant variants can occur alongside benign events, making prioritisation of important events difficult. This is especially true in cancers with genomic instability.

Transcriptome profiling has been used to interpret the functional impact of genomic variants through alterations in gene expression, transcript sequence, or both^7^. Some rearrangements that alter gene structure may be more effectively detected from RNA sequencing (RNA-seq). In particular, numerous computational approaches have been developed to reliably call fusion genes^8,9^. Similarly, a number of methods now exist to detect novel gene splicing^10–12^, and the utilisation of RNA-seq to detect and interpret splicing variants has been shown to improve diagnostic yields in rare Mendielian disorders^13,14^.

While there are many methods to detect fusion genes and novel splice variants (NSVs) from RNA-seq, transcribed structural variants (TSVs) involving a single gene, such as deletions, inversions, internal tandem duplications (ITDs) and partial tandem duplications (PTDs) are difficult to detect, and only a handful of tools have been developed^15–18^. However, these variant types can be clinically important, for example, IKZF1 deletions in Acute Lymphoblastic Leukemia^19^, FLT3 internal tandem duplications (ITDs) and MLL partial tandem duplications (PTDs) in Acute Myeloid Leukemia^15–17,20^. The prevalence of TSVs in disease is not yet known, but small SVs (<100kbp) that could give rise to TSVs are frequently seen in WGS^4,21^. In addition, fusion-finding tools are likely to miss non-canonical fusions, which we define as a fusion transcript that includes non-reference sequence (i.e. outside of the expected transcriptome) at the fusion boundary, or a fusion product created between a gene and an intergenic region.

Fusion finding tools typically use discordant or split read detection approaches, where reads are first mapped to either a reference genome^22,23^ or transcriptome^24,25^, then discordant reads are detected and a set of filtering steps are performed. While mapping information from genome alignments can be used to detect non-canonical fusions and other TSVs, most fusion finding tools do not consider these variants. However, some exceptions exist; Arriba^26^ is a method that utilises chimeric reads from STAR’s alignments to detect fusions, including non-canonical fusions, as well as PTDs. Similarly, CICERO^18^ utilises chimeric read information, and combines this with local assembly to detect canonical and non-canonical fusions, as well as ITDs. SQUID^16^, which is a method based on the idea of rearranging the reference genome using concordant and discordant reads, can detect non-canonical fusions and some TSVs. This approach has also been extended to handle heterogeneous (i.e. multi-allelic) contexts^27^. These approaches rely heavily on the reference genome and generally have limitations in detecting small variants (<100bp) such as INDELs and ITDs. These methods are biased towards detecting certain classes of variants.

An alternative to detecting variants from read alignment is to use de novo assembly. This approach attempts to reconstruct a sample’s transcriptome, then aligns the assembly back to the reference genome to identify variants. Assembly is able to reconstruct transcripts containing variants of all types and sizes, which avoids potential biases caused by alignment to the reference genome. KisSplice^28^ is an example of a reference free method focused on splice variants. It calls splice variants based on bubble-detection within the assembly De Bruijn graph. De novo assembly approaches typically generate a large number of transcripts, many of which will be misassemblies, leading to a potentially large false positive rate. Additionally, these methods are generally slower to run due to the computational requirements of de novo assembly. One strategy to reduce false positives and improve speed is to only consider a specific list of target genes, a strategy employed by TAP^15^. Another approach, used by DE-kupl^29^, is to perform differential expression of k-mers against a set of control samples. DE-kupl performs kmer counting, filtering, DE and then extends the resulting DE kmers into contigs. However, DE-kupl reports a very large number of variants (in the order of 100,000)^29^, which makes interpretation and prioritisation difficult, particularly as the tool does not provide variant type information. Another limitation of the DE-kupl approach is that it compares two groups, each with multiple samples, and is therefore not designed for detecting variants in a single sample.

Reference independent assembly combined with differential expression against controls, as introduced by DE-kupl, is a powerful strategy for the unbiased detection of transcribed variants. Here we propose an alternative approach, called MINTIE, which can detect any kind of anomalous insertion/deletion (≥7bp by default) or splicing (flanked by ≥20bp by default) in any gene. MINTIE is an RNA-seq analysis pipeline that combines the advantages of full de novo assembly with differential expression to identify unique variants in a case sample versus a set of controls. MINTIE’s pipeline includes further steps to filter, annotate and prioritise variants, which reduces transcriptional noise and aids in variant discovery.

We tested MINTIE on a simulation of 1,500 variants including fusions, TSVs and NSVs and showed that it is highly sensitive across many different variant types. We compared the performance to eight other methods and found that MINTIE was the only one able to consistently detect all classes of variants we simulated at high recall rates (>85% of total variant transcripts). In addition, we ran MINTIE on RNA-seq samples from acute myeloid leukaemia (AML) and normal blood, where we confirmed its sensitivity in detecting FLT3-ITDs, KMT2A-PTDs and fusions on real data. Normal (non-cancer) samples were used to assess the background rate of variants. In these samples, MINTIE reported a median of 122 genes per sample containing transcriptional variants, demonstrating a low background rate.

We used MINTIE to discover several interesting alterations in known driver or tumour suppressor genes in a cohort of paediatric acute lymphoblastic leukemia (ALLs) from the Royal Children’s Hospital, Melbourne. We found a recurrent non-canonical fusion involving RB1 with downstream intergenic regions of the genome. We also identified novel splicing of ETV6, and PTDs of IKZF1 and PAX5. Finally, we demonstrate that MINTIE is able to detect novel splice variants identified in a prior study in a rare disease cohort. In addition, we discovered a previously missed complex rearrangement in the disease-causing DMD gene in a patient with muscular dystrophy from this cohort.

## Results

### Algorithm overview

MINTIE is an approach for detecting novel transcribed variants that utilises de novo assembled transcripts, which are then prioritised for novelty using differential expression. The MINTIE pipeline (Figure 1) is divided into four main steps: transcriptome assembly of the case sample, pseudo-alignment of the reads from the case and controls to an index composed of the assembled and reference transcripts, differential expression to identify upregulated novel features, and annotation of novel transcripts. One case sample is required as input. The pipeline is always run in a ‘single case versus N controls’ fashion (although multiple cases versus the same control set can be run in parallel, each case is run in a 1 vs. N controls fashion). Ideally, controls should be samples of the same tissue type, but do not have to be normals or matched normals. While we recommend running MINTIE with controls, in some cases this may not be feasible, and thus the method can also be run without controls, with the caveat that more background variants are likely to be identified.

**Figure 1.**
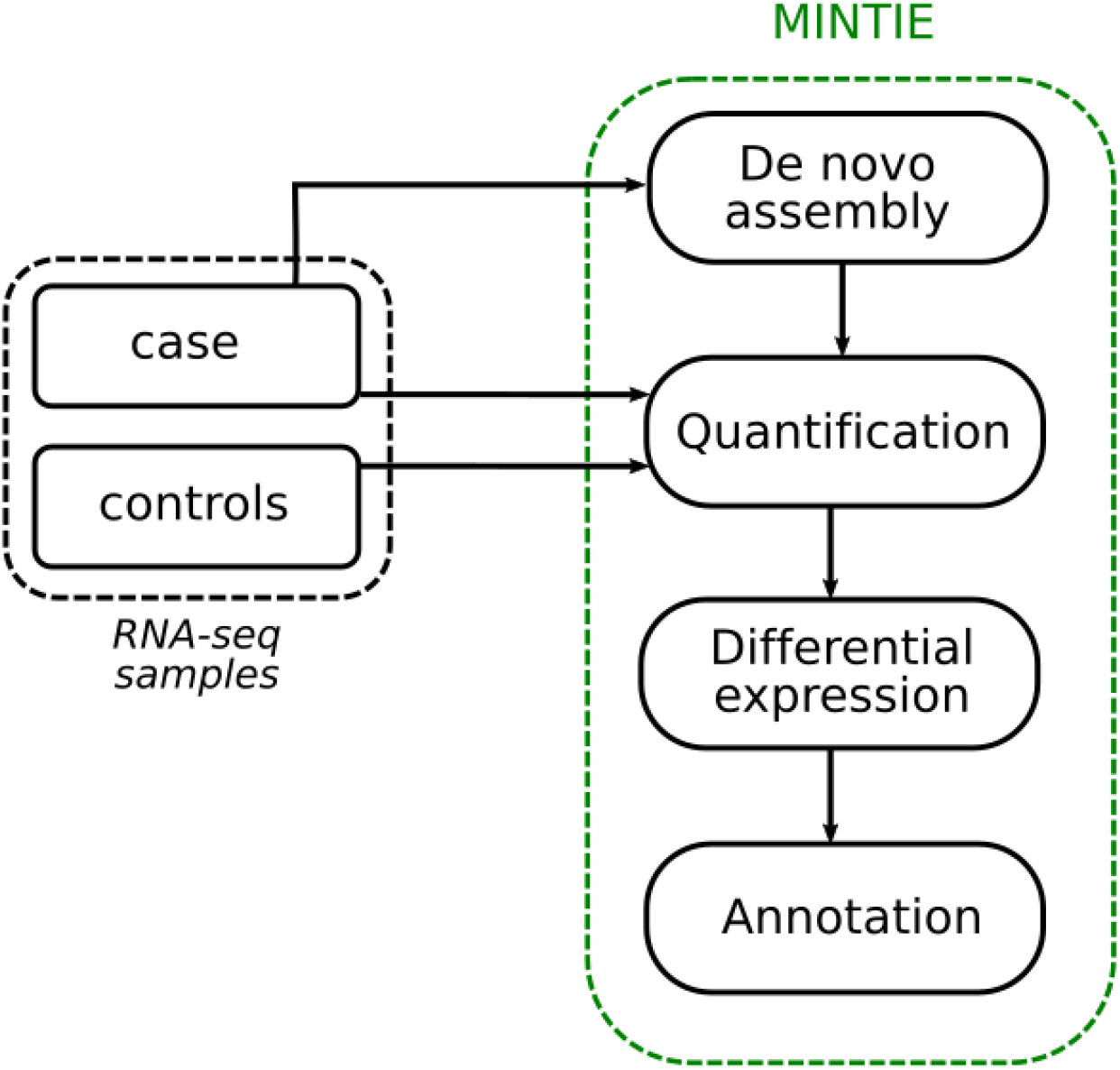
The MINTIE pipeline uses one case sample and N controls and consists of four steps: i) de novo assembly is performed on the case sample; ii) assembled *transcripts*, along with all reference transcripts, are quantified using pseudo-alignment of reads for both case and controls; iii) differential expression is performed between the case versus controls for equivalence classes that are identified as originating from assembled transcripts only, which contain novel sequence (i.e. we do not consider quantifications consistent with the reference); and iv) annotation is performed on assembled transcripts that are overexpressed in the case sample.

Conceptually, the idea of MINTIE is to perform whole-transcriptome de novo assembly, remove features consistent with reference transcripts, then perform differential expression on non-reference features versus a set of controls. Significantly over-expressed assembled transcripts are aligned to the genome and variants are identified. The novelty of this approach is that we use differential expression testing to select assembled transcripts that contain highly expressed novel sequence. Therefore, we avoid using alignment to a reference genome to define novel sequence. Instead, we identify novel sequence by testing gene expression on equivalence classes that are unique to the assembly.

In more detail, once a de novo assembly of a case sample is obtained, by default using SOAPdenovo-Trans^28^, the assembled transcripts are merged with a standard transcriptome reference (we use CHESS v2.2^29^ by default), and a Salmon^30^ index is created for this file. Salmon is then run on the case and N controls. Salmon equivalence class (EC) counts are matched across all samples. ECs represent the set of transcripts that a given read is equally compatible with, and have been used previously as a basis for differential expression^31^, differential isoform usage^32^, as well as fusion detection^33^. MINTIE retains EC counts that are compatible with de novo assembled transcripts only (no reference transcripts), and all other EC counts are discarded. This step identifies novel ECs where there is no compatible match to an existing reference transcript, as ECs containing reference transcripts are assumed to be consistent with the reference, and therefore uninteresting. The expression of these novel transcript ECs are then compared by performing differential expression testing between 1 case and N controls with edgeR^34^. This step is required to remove common but unannotated transcripts, as well as to remove erroneous assemblies where there is no discernable expression. Significant ECs (false discovery rate, FDR < 0.05 and log 2 fold change, logFC > 2 by default) are then retained, after which the transcripts corresponding to these ECs are extracted and aligned to the genome. MINTIE then performs annotation, filtering, and estimates variant allele frequency for each given variant. See Methods for more detail.

### MINTIE detects more types of variants than other tools

In order to test the utility of MINTIE across a wide range of variant types we simulated 1,500 variants by inserting, deleting, combining or duplicating sequences from 100 randomly selected hg38 UCSC RefSeq^30^ transcripts. We then generated reads from this modified reference using ART-Illumina^31^ v2.5.8. Fifteen different classes of variant were simulated (Figure 2): 500 fusions (100 each of canonical fusions, unpartnered fusions, and fusions with extended exons, novel exons and insertions at their fusion boundaries), 500 transcribed structural variants (TSVs) (100 each of deletions, insertions, ITDs, PTDs and inversions), 500 novel splice variants (NSVs) (100 each of extended, novel and truncated exons, retained introns and unannotated splice variants) and 100 unmodified background genes (see Methods). Variants were simulated at 50x coverage in a heterozygous fashion (i.e. both modified and unmodified transcripts were simulated for the variant sample, resulting in 100x total gene coverage: 50x variant coverage and 50x wild-type coverage). We also generated a control sample by simulating reads over wild-type (unmodified) transcripts at 100x for all transcripts used in the variant sample.

**Figure 2.**
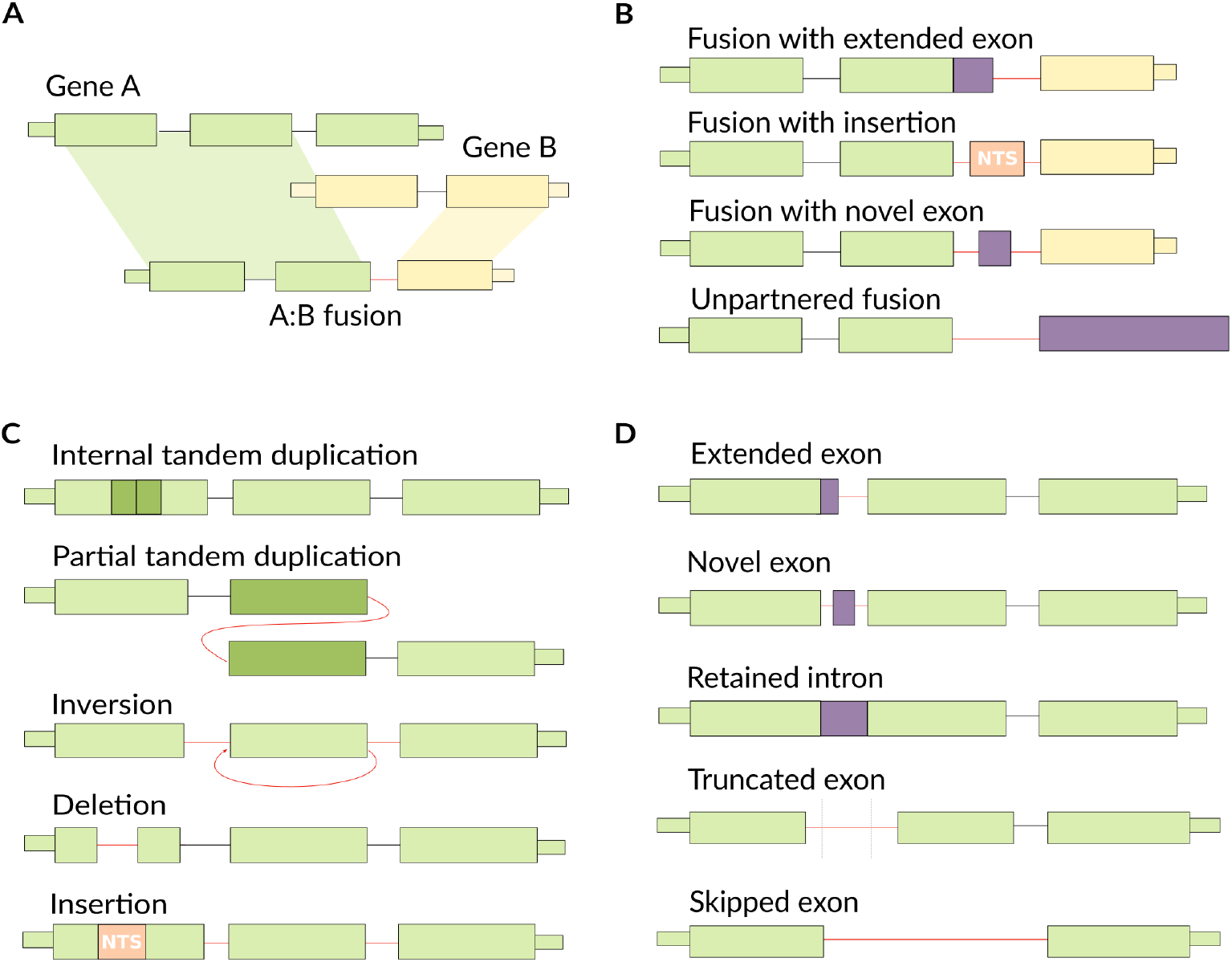
Examples of variant types detected by MINTIE. (NTS = non-templated sequence.) a. Canonical fusions. Typically defined as a transcriptional product of two genes, joined at exon-exon boundaries. b. Non-canonical fusions. Fusions that join exon-boundaries to non-gene regions, as well as fusions without a second gene partner. c. Transcribed structural variants (TSVs). May include internal tandem duplications (ITDs), partial tandem duplications (PTDs), inversions, deletions and insertions. d. Novel *splice variants* (NSVs). Includes extended exons, novel exons, retained introns, truncated exons and skipped exons.

We ran MINTIE and eight other variant detection methods (TAP^15^, Barnacle^17^, SQUID^16^, JAFFA^24^, Arriba^26^ CICERO^18^, KissSplice^11^ and StringTie^12^ on the reads generated from the simulated variants. Although DE-kupl is conceptually similar to MINTIE, we were not able to test its performance because it can not be run on a single case sample. Due to the large number of tools, each with their own output formats that had to be individually processed, we utilised a liberal approach to counting variant calls as true positives (calling a variant in a gene at either end of a fusion was counted as a hit and variant classifications were not considered; this is detailed in the Methods). MINTIE was run with control samples generated from the same transcripts without the variants (see Methods).

MINTIE detected 86.8% of fusions, 90.6% of TSVs and 79% of NSVs (Figure 3A). MINTIE had its lowest recall on unknown splice variants (skipped exons) (63%) and truncated exons (64%). Manual inspection revealed that missed cases were mostly due to the variant not being assembled or insufficient read coverage. Assembly issues could potentially be improved with another assembler. Insufficient read coverage particularly affected some deletion variants that were near the end or start of the variant transcript and thus less likely to have adequate read support. To assess the impact of read coverage we downsampled the simulated reads to three coverage values: 20x, 10x and 5x. Only a modest drop in performance was observed (Supplementary Figure 1A). Even at 5x, MINTIE found the majority of simulated variants (769 of 1,500). We also found that MINTIE retained high detection rates across all sizes of variants, even below the default insertion/deletion threshold size of ≥7bp and down to 1 bp, apart from single base-pair deletions (Supplementary Figure 2).

**Figure 3.**
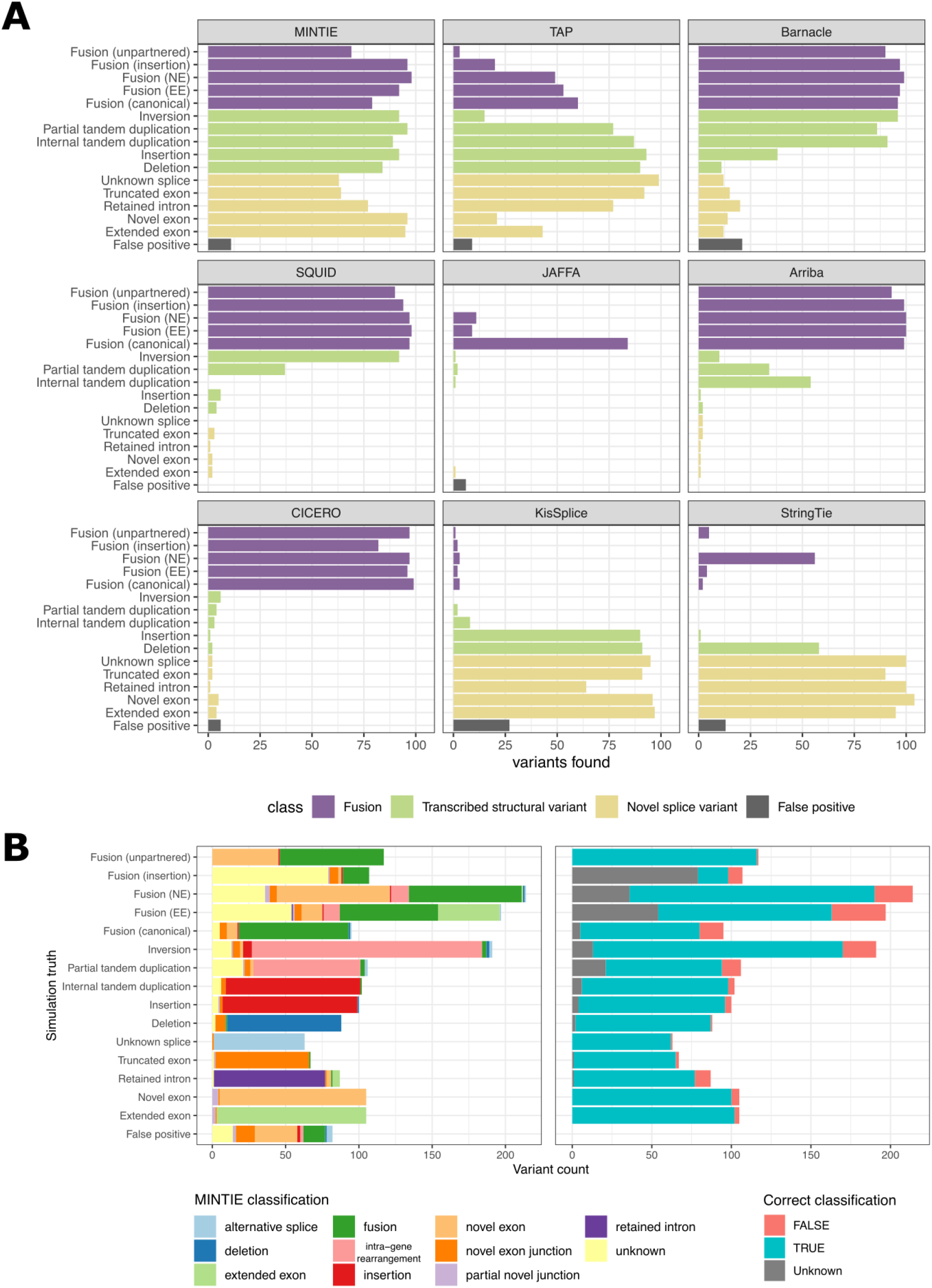
**(A)** Recall and false positives for 1,500 simulated variants for tools listed. Fusions were considered detected if either (or both) fusion partner(s) were present. **(B)** Comparison of simulated variant categories and MINTIE’s classifications for variants found in the simulated genes (left), and whether MINTIE’s classification call is consistent with the simulation (right). The ‘unknown’ classification refers to soft-clip variants, which could indicate a number of types. MINTIE may call multiple variants per event (e.g. fusion + novel exon), so some categories have more than 100 events.

While MINTIE had lower sensitivity when compared to specialised tools for novel splice variant detection (StringTie and KisSplice), our method was able to find the most total variants (86.2%), while no other tool came close to this. CICERO has been reported to have high sensitivity for ITD detection^18^, however it only identified 3 ITDs in our simulated data set. CICERO ranks ITDs by whether the gene is known for recurrent ITDs. We did not simulate ITDs in such genes and hypothesise that this is the reason they were missed. These ITDs were reported (93%) in CICERO’s unfiltered output (Supplementary Figure 3), however the unfiltered output also included a large number of false positives (596 in total). Given that only 100 unmodified (wild-type) transcripts were simulated, we expected minimal false positives to be identified. No false positive calls were identified for SQUID and Arriba. JAFFA reported six false positive fusion genes (across five calls, all at low confidence). CICERO, TAP, MINTIE, StringTie and Barnacle called 6, 9, 11, 13 and 21 false positives respectively, while KisSplice had the highest number at 27. StringTie, TAP and Barnacle reported calls in background genes (those simulated without any variants) with 1, 2 and 3 hits respectively. MINTIE, KisSplice and JAFFA did not report any false positive calls in background genes. All other reported false positives were outside of simulated gene regions. Manual inspection of MINTIE’s false positives revealed that these calls were largely due to poor alignment to homologous sequences (for example, to a pseudogene of the variant gene).

To aid with variant interpretation and prioritisation, MINTIE outputs classifications for the type of transcribed variant found: fusion, intra-genic rearrangement, deletion, insertion, novel/extended exon, novel exon junction and retained intron. We explored the classification accuracy of MINTIE’s results using the simulation. Figure 3B shows that the variant types MINTIE assigned to variants called within the simulated genes are mostly correct (86.5%) across the 15 categories.

### MINTIE identifies known fusions and ITDs in Leucegene samples

In order to demonstrate the sensitivity of MINTIE in real samples, we applied MINTIE to a set of 77 AML samples with known fusions, internal tandem duplications and partial tandem duplications (Supplementary Note 1). This included the NUP98-NSD1 fusions that were validated from Lavelle et al.^33^ (N=7). We also tested MINTIE on samples containing CBFB-MYH11 fusions, RUNX1-RUNX1T1 fusions and FLT3-ITDs known to be in the core binding factor (CBF) AML data^27^, as well as a cohort containing KMT2A-PTDs^32^, identified by Audemard et al.^33^. Although we benchmarked just five variants across three variant types, which represents a small subset of the types of events MINTIE can detect, it enabled MINTIE’s sensitivity to be confirmed using well known and validated events. The Leucegene data was also used to explore the impact of control choice, read coverage in real data and rate of background events.

In all these analyses, we used 13 normal Leucegene samples as controls (5 granulocytes, 5 monocytes and 3 total white blood cells). In the AML data sets, a median of 592 variant genes were reported per sample (range 261-2265). MINTIE detected 50/53 total fusions (Figure 4A); manual inspection revealed that two CBFB-MYH11 fusions were missed due to low expression and one NUP98-NSD1 fusion was not assembled. Additionally, all FLT3-ITDs were detected in the CBF and NUP98-NSD1 cohorts (default parameters needed to be adjusted for one patient sample in order to detect a small 3bp FLT3-ITD). MINTIE was able to find 16/24 KMT2A-PTDs in the samples identified by Audemard et al.^33^ when using normal samples as controls and 19/24 when using no controls (Supplementary Table 1). Some of the variants that were successfully detected (using normals as controls) had estimated expressed variant allele frequencies (VAFs) down to 0.0580 (median 0.2220, max 0.7127), suggesting that only a small amount of the total gene expression needs to come from the variant isoform for detection. The variants that were missed had low coverage (<17 reads). Of the 8 missed variants, 6 were due to insufficient assembly and 2 were filtered out by the counts per million (CPM) filter due to low counts in their respective ECs.

**Figure 4.**
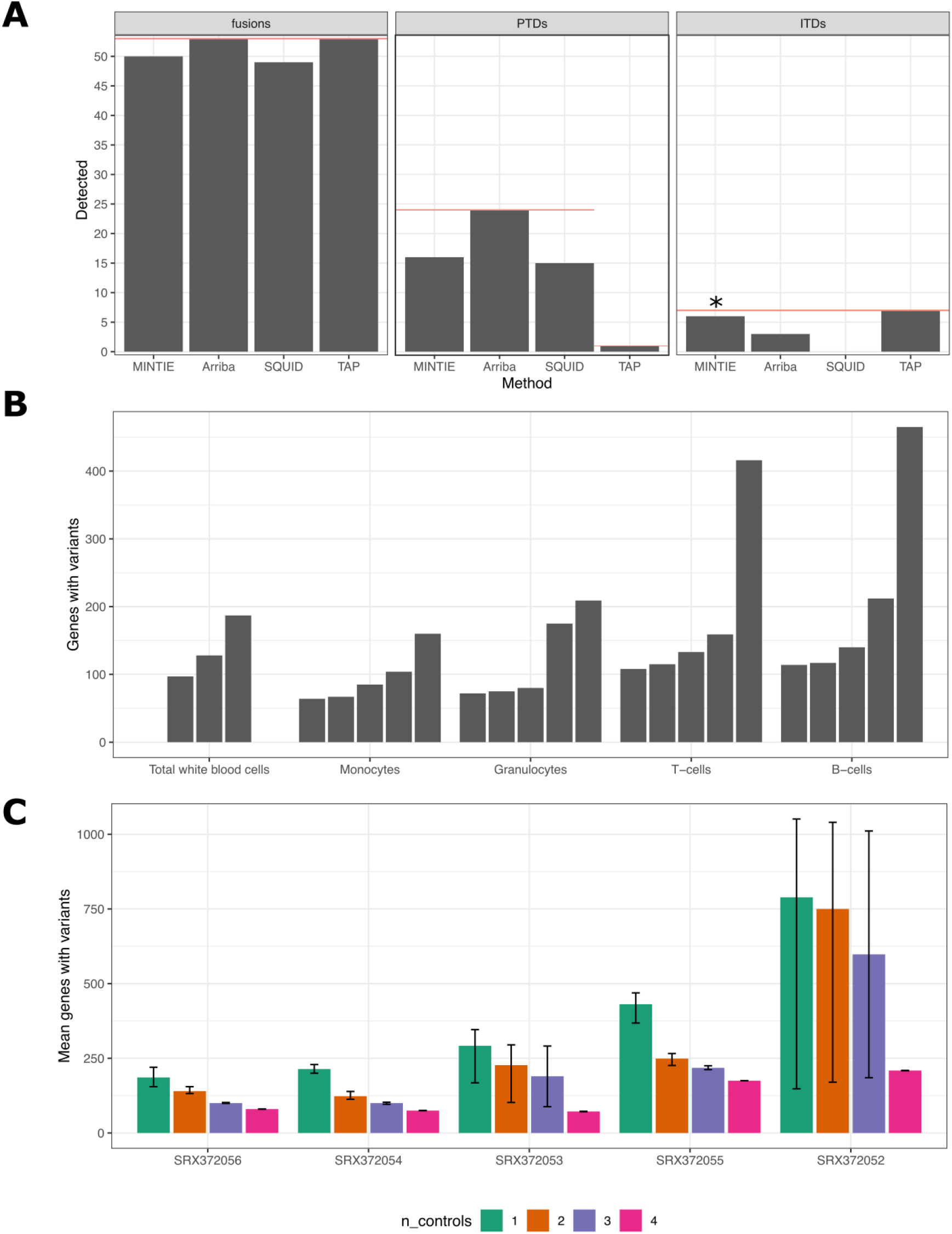
**(A)** Summary of true positive variants detected by MINTIE, Arriba, Squid and TAP in Leucegene AML samples. *Red line shows the total variants present in the category* (only results from one sample were available for TAP from the KMT2A cohort). * The remaining 3bp FLT3-ITD could be detected by changing MINTIE’s minimum gap size parameter to 3. **(B)** The number of genes with variants found per sample across the 23 non-cancer cell types from Leucegene. Each bar represents a different sample within the cell type. Samples are ordered by the total number of genes with detected variants. **(C)** Mean number of genes with variants called in Leucegene granulocyte samples where different numbers of (granulocyte sample) controls were used. ‘Error’ bars show minimum and maximum variant genes called across replicate experiments, reflecting different combinations of controls. N = 4 has one combination of controls, N = 3 has two, N = 2 has three and N = 1 has four.

We also analysed these Leucegene samples with three selected state-of-the-art methods, which are designed for these variant types: Arriba (a representative fusion finder), SQUID (a representative large TSV caller) and TAP (a general purpose caller). Arriba found all fusions and KMT2A-PTDs, but missed more than half the FLT3-ITDs (Figure 4A). Squid performed similarly to MINTIE, finding 49/53 fusions and 15/24 KMT2A-PTDs, but failed to identify any of the FLT3-ITDs. This highlights how current tools designed to detect larger rearrangements such as fusions and PTD may be insensitive to smaller events such as ITDs. TAP was able to detect all events (Figure 4A). However, due to difficulties in installing and running TAP, detection rates were taken from their publication^15^, and results from only one KMT2A-PTD sample were available. These results show that MINTIE has sensitivity within the range of existing state-of-the-art tools designed specifically for detecting these known, well defined variants. The validation cohort incorporates only three of the 15 variant types from our simulation study and the PTDs and ITDs were focused on a single gene in each case (KMT2A and FLT3 respectively). Thus, they represent only a fraction of MINTIE’s utility. Nevertheless, these results are consistent with our simulation and suggest that MINTIE is likely to have good sensitivity over a range of events.

### MINTIE detects a low number of background variants

We applied MINTIE to a set of 23 non-cancer healthy adult sorted blood cell samples obtained from Leucegene^34^. We expected these samples to have low numbers of transcriptional variants compared to cancer samples, and thus provide an estimate to the background rate of detected variants found by MINTIE. This set was composed of several cell types: T cells (N=5), B cells (N=5), granulocytes (N=5), monocytes (N=5) and total white blood cells (N=3). Each sample was run with MINTIE using all other samples of the same cell type as controls.

Figure 4B shows the total number of genes containing transcriptional variants identified across the non-cancer samples. In general, MINTIE had a low background variant rate, with a median of 122 affected genes across samples (range 61-1,397). We identified 19,893 expressed genes across the non-cancer samples (>1 CPM in at least one sample), thus the range translates to a median of 0.6% of expressed genes identified as containing variants. 83.3% of the 2,363 identified variant genes were protein coding (compared to 74.39% of expressed genes that were protein coding). 56.2% of variants were classified as novel splicing variants and may represent true variation in the samples. The number of variants called was variable between samples, even within samples of the same cell type. We found only a weak correlation between library size and number of transcriptional variants detected (spearman correlation coefficient = 0.2105). A moderate fraction of variants (23.3%) were of unknown type and may be due to poor alignment of the de novo assembled transcripts to the genome. We note that one B-cell sample contained the largest number of variant calls (1,710 variants). Upon manual inspection, we noted that 1,259 of these variants were clustered in a 1MB region at the end of the long arm of chromosome 14 containing immunoglobulin genes, likely indicating that these genes, which are commonly rearranged, were highly expressed in this sample.

### Well-matched controls reduce the number of background variants

We found that using correctly matched controls was important for filtering out the normal range of transcriptional diversity in a given sample. We compared the number of variant genes observed in normal total white blood cells (TWBCs) when using different controls: the same cell type (TWBCs), as well as granulocytes, monocytes, T-cells and B-cells. Using the other TWBCs as controls resulted in the fewest number of variant genes found across all three samples (N=412), followed by monocytes (N=715) and granulocytes (N=763) (Supplementary Figure 4). This corresponds with the expression similarity (Supplementary Figure 5). B-cells and T-cells cluster together in gene expression, distinct from TWBCs, and using them as controls identified the most variant genes at 1,564 and 1,630 respectively. These results confirm that comparing against controls of similar expression profiles combined with enough control samples resulted in fewer total number of variants found.

To further explore the effect of control number and similarity on MINTIE’s variant calls, we performed an experiment using 1-4 controls in different combinations using the Leucegene granulocyte samples. We observed fewer variant genes called as the number of controls were increased (Figure 4C). One sample (SRX372052) displayed notably higher variability in variant genes called, compared to other samples, when using 1-3 controls. On inspection of expression similarity through a PCA plot (not shown), the sample was revealed to cluster more closely with two samples than the others. When the less similar samples were randomly chosen as controls, variant numbers were significantly higher. These results suggest that using more controls reduces the number of background variants called and confirms that sample similarity will impact the number of variants called.

We also ran the cohort of 24 Leucegene AML samples containing KMT2A-PTDs against three different sets of controls: i) 13 normal (non-cancer) controls (5 granulocytes, 5 monocytes and 3 total white blood cells), ii) a reduced set of 3 normal controls containing one sample from each cell type, iii) a set of 13 AML samples from a different cohort, and iv) no controls. MINTIE was able to detect 19/24 variants using no controls, 16/24 variants using the normal controls, 15/24 variants using the reduced normal set, and 11/24 variants using AML controls (Supplementary Table 1). The lower detection rate versus other cancers was due to high read counts in the controls for the ECs corresponding to the KMT2A variant. While there was no evidence of KMT2A rearrangements in the control samples used, some of the assembled contigs contained other, more common variants in the same EC (such as an extra A insertion in a small A repeat region proximal to the breakpoint), and thus were not found to be significant at the DE step. Differences were observed in total variants found using different control sets (Supplementary Figure 6), with a median of 645.5 variant genes (range 264-2265) found per sample for the 13 normal controls compared to 913 median variant genes (range 490-2552) for the reduced set of 3 non-cancer control samples and a median of 211.5 variant genes (range 130-2852) found when using the 13 AML controls. Using no controls yielded significantly higher variant numbers (range 4750-11796, median 9255.5), indicating that using even small numbers of controls acts as an effective filtering strategy. If we assume most variants are background events, unrelated to cancer, this suggests that using samples with well matched transcriptomes as controls reduces the number of detected background variants.

### MINTIE identifies novel transcriptomic variants in B-ALL

In order to test MINTIE’s ability to discover previously undetected events we applied MINTIE to a set of 91 samples across a cohort of 87 paediatric B-ALL patients from the Royal Children’s Hospital, Melbourne Australia^35^. As the control group we used a subset of 34 samples from the cohort with already known driver fusions, identified by molecular testing, and validated in the RNA-seq with fusion calling (Supplementary Note 2). A median of 48 variant genes was found per sample (range 15-633, Supplementary Figure 7). We filtered on variants found in 379 recurrently altered genes (not considering immunoglobulin genes) in over 2,500 paediatric cancer samples reported in two landscape papers^36,37^, and found 339 variants across 131 of these genes. The top recurrent genes were ETV6 (26 variants across 8 samples), IKZF1 (14 variants across 7 samples) and IKZF2 (13 variants across 3 samples) (Supplementary Figure 8).

One sample contained an in-frame, single-exon tandem duplication in IKZF1. In-frame deletions of IKZF1 are recurrent in paediatric ALL and associated with poor prognosis^38^. They are believed to act in a dominant negative manner and we hypothesise that this might also be the case for the IKZF1-PTD we detected. We also found one sample with an in-frame tandem duplication of 4 exons in PAX5, which correlates with previously reported amplification of PAX5 exons in paediatric ALL^39^. A subtype of B-ALL is defined by PAX5 alterations, with a range of diverse variant types across patients (fusions, amplifications and mutations)^40^.

In addition we found three samples with exon skipping and cryptic exons of ETV6 (Figure 5A). In one sample, Case 1, we observed a unique alternative splicing event between exons 2-5, resulting in lowered expression of intervening exons, indicating a potential deletion. Exon skipping due to an intragenic deletion in EVT6 has previously been reported in paediatric ALL^41^. Karyotyping of the sample indicated a complex rearrangement of ETV6, but no fusion involving ETV6 was found. The same exon skipping event was observed in a single TCGA Lung Squamous Cell Carcinoma sample, found with SeqOthello^42^. In the other two B-ALL samples, Case 2, which were from diagnosis and relapse of the same patient, we observed two novel ETV6 isoforms: splicing between exons 2-8 and splicing between exon 5 and two downstream novel intergenic exons. Expression of exons 6 and 7 was absent.

**Figure 5.**
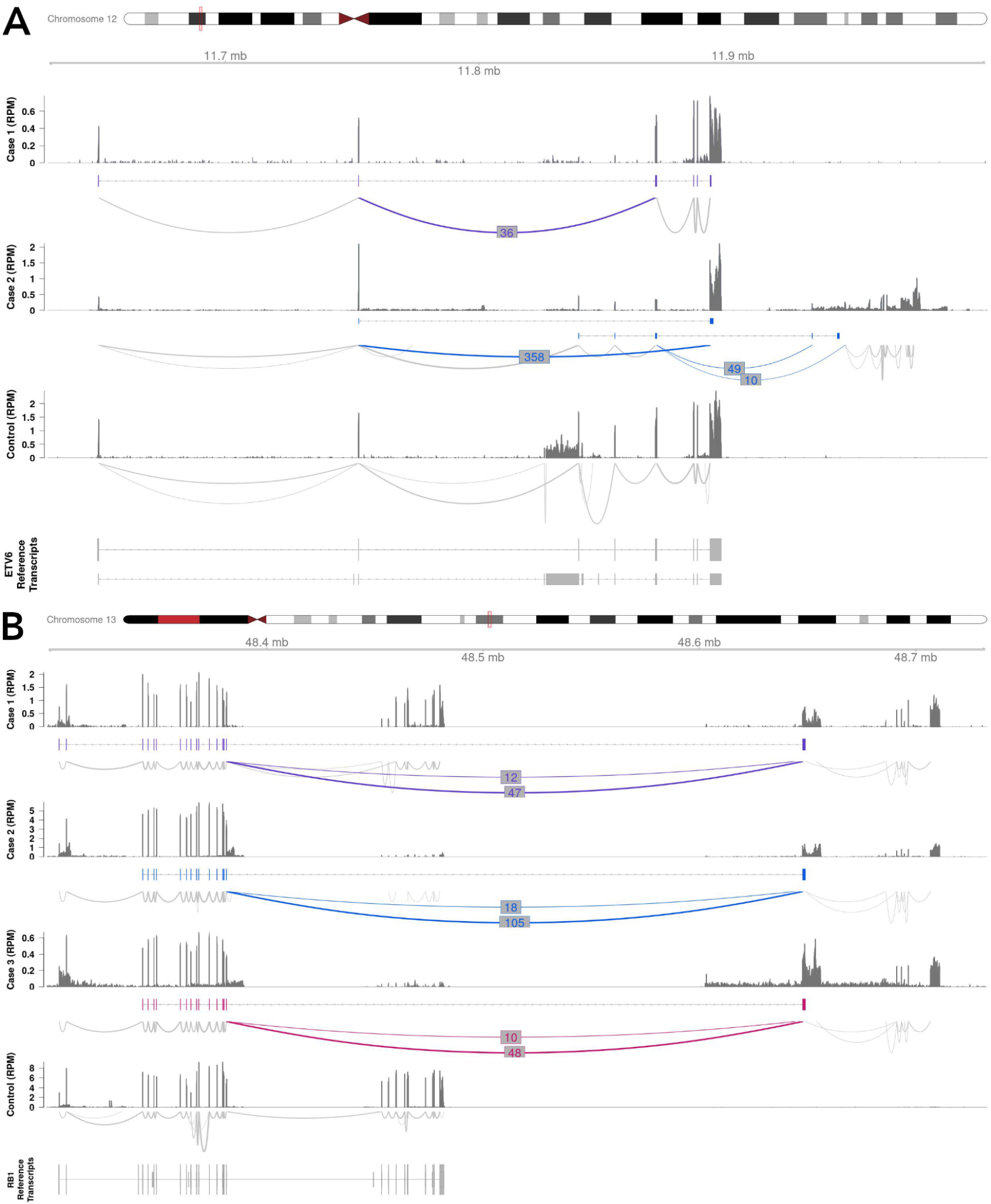
Examples of novel variants detected in a cohort of paediatric B-ALL patient samples. **(A)** Two cases with altered splicing of EVT6, and **(B)** three cases with RB1 unpartnered fusions. Shown for each case in top to bottom order is the read coverage, novel assembled transcript and splicing across the variant. Novel splice junctions are coloured with their corresponding number of supporting reads. There are two acceptor sites (right side) for the RB1 variant, approximately 3.5kb apart, that correspond to alternative splice variants; the donor site (left side) is always the same. For comparison, a representative control sample is shown (bottom) with corresponding expressed reference transcripts.

A B-ALL subtype classifier based on gene expression (https://github.com/Oshlack/ALLSortsv0.1) was run on the four diagnosis samples harbouring ETV6 splicing, IKZF1-PTD and PAX5-PTD variants. All samples were found to have a strong probability of belonging to their respective class, ETV6-RUNX1-like, IKZF1 N159Y and PAX5alt (data not shown), suggesting these events may be drivers.

MINTIE also detected an unpartnered recurrent fusion involving the tumour suppressor gene RB1 from the end of exon 17 to an intergenic region approximately 165kb downstream (Figure 5B) in three other samples. In all three samples, two splice variants of the fusion were seen, which involved different novel downstream exons (Supplementary Table 2). Read coverage dropout suggested a 188kb genomic deletion involving the 3’ end of RB1 in one sample, however, this was not clear in the two other samples. RB1 focal deletions have been reported in both B-ALL and T-ALL^37^ from DNA-based assays, supporting this hypothesis. We further expanded our search for this variant by querying TCGA using SeqOthello^42^. One of the fusion splice variants involving RB1’s exon 17 and the same downstream intergenic region was detected in two other samples, a breast invasive carcinoma and an esophageal carcinoma. MINTIE successfully detected the variant in both of these samples when we obtained the RNA-seq data and processed it through the pipeline, using 3 normal controls from TGA. These results suggest that focal RB1 deletions occur in multiple cancer types, and form a stable transcript that can be detected from RNA-seq alone using MINTIE.

### MINTIE identifies novel splicing and transcribed structural variants in rare disease

To demonstrate the utility of MINTIE in rare disease, we analysed RNA sequencing data from patients with rare muscle disorders from Cummings et al.^14^. In the study, diagnosis was made by identifying altered transcripts using RNA-seq in pathogenic genes with an identified variant of unknown significance. From this data set there were 10 patients that had RNA-seq available with 13 splicing variants that could potentially be detected by our method. We used 10 muscle RNA-seq samples from GTEx^43^ as controls, and ran MINTIE transcriptome wide using RNA-seq alone without knowledge of which genes had DNA mutations. MINTIE identified the correct novel splice variants for 9/13 events (Table 1).

**Table 1.** List of variants detectable by MINTIE from Cummings et al.^14^ rare muscle disease data.

| Patient | Gene | Junction 1 | Junction 2 | DNA variant | RNA variant | Detected Variant |
| --- | --- | --- | --- | --- | --- | --- |
| E2 | NEB | chr2:151687733-151690727 |  | SNP | skipped exon | Y |
|  | NEB | chr2:151687733-151688175 |  | SNP | extended exon | Y |
|  | NEB | chr2:151687733-151688007 |  | SNP | extended exon | Y |
| C9 | NEB | chr2:151531896-151533330 |  | SNP | extended exon | Y |
| N25 | NEB | chr2:151498352-151498491 |  | SNP | novel exon | Y |
| N31 | COL6A1 | chr21:45989778-45989894 | chr21:45989965-45990258 | SNP | novel exon | Y |
| N32 | COL6A1 | chr21:45989778-45989894 | chr21:45989965-45990258 | SNP | novel exon | Y |
| C11 | RYR1 | chr19:38467720-38468966 |  | SNP | truncated exon | Y |
| C1 | POMGNT1 | chr1:46194651-46195811 |  | SNP | skipped exon | N (low CPM) |
|  | POMGNT1 | chr1:46189357-46189458 |  | SNP | retained intron | Y |
| E4 | TTN | chr2:178580609-178581906 | chr2:151498552-151499298 | INDEL | skipped exon | N (not assembled) |
| N22 | TTN | chr2:178777317-178777460 |  | SNP | truncated exon | N (low CPM) |
| C3 | DMD | chrX:31729748-31819975 |  | inversion-deletion | skipped exon | N (EC not significant) |

Two variants were filtered out due to low expression (exon skipping in POMGNT1 and exon truncation in TTN), while an exon skipping event in TTN which was only supported by 2 junction reads, failed to be assembled. A transcript harbouring the DMD skipped exon in patient C3 was not found to be significant in the DE step, as a sizable number of reads supporting the variant transcript were found in the controls. Manual inspection revealed that the assembled transcript for this DMD case contained two variants, one being the target variant (the skipped exon), and a secondary variant (a homozygous SNP). The secondary variant was not in the reference transcriptome but was found in 9/10 controls. Cases such as this are rare, but represent one potential weakness of an EC-based method, whereby multiple variants assembled into the same transcript cannot be resolved at the EC-level.

Patient samples C2 and C4 were identified with large-scale inversions in the DMD gene using DNA sequencing that manifested as lower exonic coverage across regions of the gene. Cummings et al. did not report an exon skipping RNA-seq feature, and thus we could not check specific boundaries in the RNA-seq. However MINTIE did find novel transcriptomic variants in DMD in both of these samples. Running BLAT^44^ on these soft-clip sequences revealed that they aligned on the opposite strand to the gene sequence, supporting an inversion variant. This is an example where MINTIE can provide insight into DNA-level structural variants solely from the RNA-seq.

We also investigated whether MINTIE could detect any variants of interest that were missed by the Cummings et al. study. Prioritising the same set of NMD candidate genes as the study’s authors, and stringent filtering on MINTIE’s outputs, together with manual validation (see Methods), we identified five variants: four deletions and a fusion across eight patients (Supplementary Table 3). All deletions were found to be rare UTR variants found in the general population (<5% MAF), of which, two were reported in ClinVar (DMD’s 3’ UTR deletion and VAPB’s 3’ UTR deletion) as variants of unknown clinical significance. The VAPB mutation was reported to be associated with Spinal Muscular Atrophy (ClinVar Submission Accession SCV000434613). Most interestingly, we identified a fusion transcript in DMD in a patient where exome sequencing suggested that DMD was a strong candidate gene^14^, however, no corresponding RNA variant was found in the study. The unpartnered fusion joined the end of exon 43 of DMD on the X chromosome to an intergenic region on chromosome 8 with two cryptic exons (Figure 6). Upon manual inspection, we noted 10 split-reads with three different soft-clip sequences at the exon 43 boundary; of these, 5 sequences corresponded to the breakpoint on chromosome 8, matching the assembly as expected. However, two other sequences were found to correspond to regions 650bp upstream (2 split-reads) and 70kb downstream (3 split-reads), suggesting multiple potential fusion transcripts. We confirmed the presence of this rearrangement with the original authors through private correspondence, who independently discovered it using whole genome sequencing and recently reported it in Waddell et al, 2021^45^.

**Figure 6.**
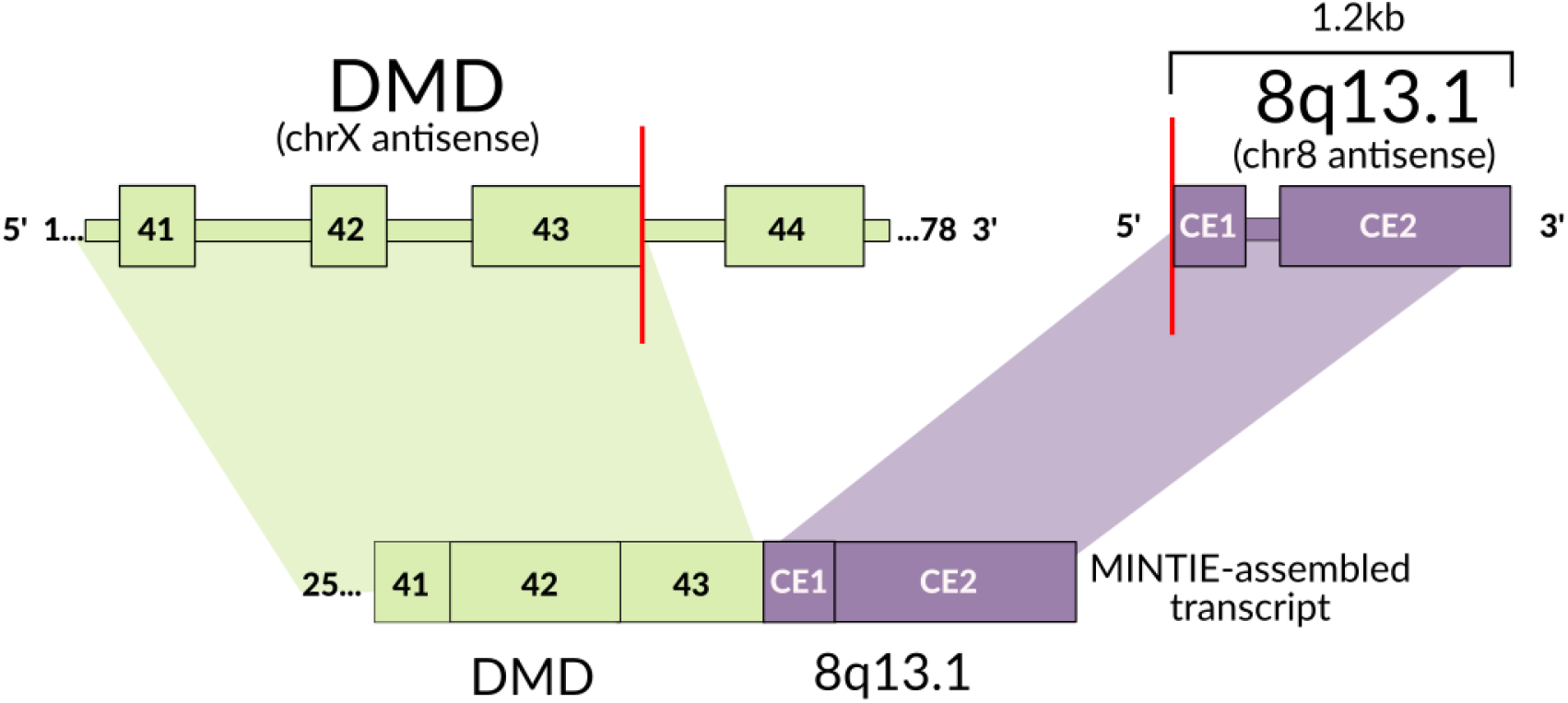
Unpartnered DMD fusion found in a rare disease sample (CE = cryptic exon). MINTIE’s assembled transcript indicated a fusion boundary at DMD’s exon 43 to an intergenic region in chromosome 8.

### Computational Performance

Computer resources are often seen as a limitation to the application of de novo assembly in variant detection. In MINTIE, we used SOAPdenovo-Trans as the default assembler due to its computational efficiency compared to other assemblers. MINTIE also leverages pseudo-alignment to minimise compute times. To benchmark performance, we selected three samples (containing the minimum, median and maximum variants called) from the Leucegene KMT2A-PTD cohort and ran MINTIE against the set of 13 controls previously described (Table 2). Tests were run on a server with 32 cores and 190 GB of available memory with MINTIE’s bpipe pipeline concurrency limited to 32. The variant numbers also corresponded with the number of read pairs, ranging from 67 million to 113 million. Compute times were between 3 hours and 24 minutes and 7 hours and 4 minutes, where the run time scaled with the number of reads. Maximum memory usage did not appear to be significantly correlated with the read number, as all samples were between roughly 90-100 GB max memory usage (we note that some samples may utilise >190 GB memory, primarily due to the assembly step). The first three steps, dedupe, trim and assemble steps account for approximately over half the run time, and most of the memory usage. For instance, in one sample these stages took 4 hours and 4 minutes and used, at maximum, 102.64GB of memory. The rest of the pipeline finished in 3 hours and used at maximum 16.95 GB of memory. These benchmarking results demonstrate that MINTIE’s utility is not hindered by computational considerations, and a large number of samples can be processed on a medium sized server in a reasonable time frame.

**Table 2.** Compute times for three selected samples from the Leucegene KMT2A-PTD cohort, including the sample with the least variants called, the median, and the most. Reads were 100bp and samples were run against 13 AML controls on a server with 32 cores and 190 GB of available memory.

| Sample | Variants | Read pairs | Time | Max mem (GB) |
| --- | --- | --- | --- | --- |
| 08H012 | 128 (min) | 67,789,336 | 3h24m | 101.47 |
| 07H155 | 214 (median) | 98,946,255 | 6h42m | 89.85 |
| 10H007 | 2093 (max) | 113,017,564 | 7h04m | 102.64 |

## Discussion

Here we present the MINTIE pipeline, a caller for detecting a broad range of transcribed variants ≥ 7bp in size. MINTIE excels at detecting unusual and rare upregulated variants. MINTIE frames variant calling as a problem of detecting sample specific expression of transcripts with novel sequence, without relying on read alignment to the reference genome. MINTIE achieves this by performing de novo assembly to reconstruct all transcripts expressed in a sample. It then leverages the unique approach of using equivalence class counts to reliably detect differential expression of unannotated sequence compared to a set of controls. Once novel transcripts are identified, MINTIE performs a number of steps to filter out transcripts found in the reference annotation and finally annotates the remaining variants into subtypes.

We demonstrated MINTIE’s ability to find a diverse range of fusions, structural and splice transcriptomic variants by simulating 1,500 variants, where MINTIE achieved >86% recall. We further benchmarked fusion callers, transcribed structural variant callers and splice assemblers and found that MINTIE detected the full range of simulated variant types, while the other methods could not. We validated the ability of MINTIE to detect fusions and duplications in a set of validated AML cancer samples. Even for fusion finding, where many reliable methods exist, MINTIE’s sensitivity was comparable. We propose that MINTIE could serve as either a first-pass scanner that is able to detect a large range of variants including fusions, or as a transcriptome-wide discovery tool for rare variants if nothing is found using conventional fusion finders.

Using the Leucegene data, we showed that using several well matched controls are important for reducing the number of background variants. For many cases, other disease samples with similar expression profiles can be used as controls. In this instance we advise using samples with known driver variants, so that the unique variants to be detected are unlikely to be present in the controls. Identical variants that are present in the set of controls will limit detection sensitivity in the case sample.

MINTIE has the potential to discover a wide range of important events that have thus far evaded detection due to the limitation of conventional analysis approaches. We demonstrated this by running MINTIE on a cohort of 87 B-ALL samples, where we found interesting events in disease relevant genes, including 3 samples with the same RB1 unpartnered fusion, IKZF1 and PAX5 PTDs, and 2 samples with altered splicing of ETV6. On rare disease samples, we showed that MINTIE could detect novel splice variants, and importantly, was able to discover diagnostically relevant genomic rearrangements from RNA-seq alone.

It is important to consider some of the limitations of MINTIE. The methodology has been optimised to detect a large number of variant types and thus some specialised approaches geared at detecting fewer variant types outperform MINTIE in accuracy and runtime. MINTIE also requires sufficient expression of the variant relative to the controls. A further consideration is that MINTIE is, to some extent, reliant on appropriate control samples. MINTIE is a method based on RNA sequencing, and can only identify structural rearrangements that affect transcribed regions. Similarly, lowly expressed genes or variants that knock out the expression of a gene will be difficult to detect. Finally, MINTIE utilises de novo transcriptome assembly to reconstruct transcript sequences. While this has the advantage of allowing complex variants to be detected, the run-time and RAM requirements are generally higher than many of the alignment based approaches and the shortcomings of this approach also affect the method, such as difficulty with repetitive regions. We chose an assembler with good performance while minimising compute time. Slower, more computationally intensive assemblers may reconstruct more accurate transcripts. Although we have not benchmarked the effects of different transcript assemblers, MINTIE allows an assembled transcriptome from any source to be passed to the program bypassing default assembly by SOAPdenovo-Trans. Looking forward, long read transcriptome sequencing technology is readily being adopted and MINTIE could be used with polished long read data in tandem with short reads, avoiding the disadvantages of assembly.

Here we have presented a novel, fast, transcriptome-wide approach for detecting rare transcribed variants, and have demonstrated the use of the tool through simulations, real cancer and non-cancer samples and have demonstrated the detection of a number of alterations, some of which are clinically relevant. We propose that MINTIE will be used to identify novel drivers and loss-of-function variants in cancer and in rare diseases. Unlike existing methods, which are targeted at specific types of rearrangements or splicing, or specific genes, MINTIE is more agnostic to variant type. We have tested MINTIE on several types of variants, but there are potentially many other types which MINTIE could detect. For example, retrotransposon insertions into genes or complex events involving multiple rearrangements, which could be generated by chromoplexy or chromothripsis. Little is known about the frequency or characteristics of transcribed variants, but we now have a method to uncover and study them.

## Methods

### Pipeline detail

#### De novo assembly

RNA-seq reads of the case sample are de-duplicated using fastuniq^46^ v1.1 and trimmed using Trimmomatic^47^ v0.39 and then de novo assembled using SOAPdenovo-Trans^48^ v1.03 with kmers of 29, 49 and 69 (by default; in all samples presented in the paper, we used kmer sizes 49 and 79 unless otherwise indicated). (Optionally, Trinity^49^ or rna-SPAdes^50^ may be used for assembly, or the user may provide their own assembly.) The different kmer assemblies are merged, sequences are deduplicated and assembled transcripts longer than 150 bp (by default) are retained.

#### Quantification

all assembled transcripts are merged with the CHESS^51^ v2.2 transcriptome reference sequence, which is then indexed by Salmon^52^ v0.14. Salmon is then run on this index for the case and control samples in single-end mode (otherwise short assembled variants may not be correctly counted). Sequence bias is corrected (--seqBias), equivalence class (EC) counts are extracted (--dumpEq), mapping validation (--validateMappings) and hard filtering (--hardFilter) are enabled.

#### Differential expression

ECs are matched between all samples by membership (i.e. an EC composed of transcripts A, B and C in sample 1 is given an identifier, and will be considered the same EC as one composed of the same transcripts in sample 2). Library sizes are calculated prior to any filtering. ECs containing *only* assembled transcripts (i.e. the EC does not contain any transcripts from the reference) are kept and, after light filtering (CPM > 0.1 in the case sample), the case sample is compared to the controls using edgeR^53^ v3.26.5. Dispersions are estimated using the classic (non-GLM) model^54^, and transcripts are fitted using edgeR’s GLM quasi-likelihood fit function^55^ with the robust flag set to true^56^. edgeR’s GLM likelihood-ratio test^57^ is used to perform differential expression. All transcripts associated with significant ECs are retained (FDR < 0.05 and logFC > 2 by default; we used a logFC > 5 for all analyses unless otherwise indicated).

#### Annotation

all significant transcripts are aligned using GMAP^58^ (2020-06-04) to the hg38 genome with all alternative reference contigs removed. GMAP’s `--max-intronlength-ends` parameter is set to 500kb to prevent soft-clipping at long terminal introns, and `--chimera-margin` is set to correspond to MINTIE’s clip length (default = 20). All aligned transcripts are extracted and compared to the CHESS^51^ v2.2 reference GTF to identify fusions, TSVs and novel splicing. We require that at least 30 base-pairs and 30% of the transcript sequence is aligned somewhere in the genome to retain a candidate transcript. Transcripts not overlapping any known reference exons are discarded. Broadly, we set the criteria for novel variants as: i) the transcript contains an insertion or deletion of at least 7 base-pairs, ii) the transcript has a soft-clip or hard-clip (based on the GMAP’s alignment) that is least 20 base-pairs in length, iii) the transcript contains a splice junction not seen in the reference transcriptome, and iv) the transcript contains a block of at least 20 base-pairs that is not normally transcribed in the reference genome. Passing category (i), (ii), (iii) and/or (iv) will retain the transcript as potentially harbouring a candidate variant. We discard any variants matching category (iv) where no splicing occurs (i.e. the transcript is aligned as an unspliced block) unless the transcript’s start and end are contained within flanking exons, indicating a potential retained intron.

Annotated variants are further refined by matching novel blocks with novel splice junctions and identifying valid donor/acceptor sites for novel junctions (defined as observing the sequences AG/GT or AC/CT before the splice junction). By default, the motif checking allows tolerance for one mutation on either end of a junction (this can be made more or less lenient by the user). As extended/novel exons should create novel exon boundaries, we consolidate these variants and discard any extended/novel exons not supported by a corresponding novel junction. Novel and extended exons must be accompanied by at least one novel junction at either (or both) ends of the block and have a valid donor or acceptor motif. Alignment gaps (of at least 7bp) within single exons are checked to capture potential novel intron, and exon ends are checked for truncation by at least 20bp. A single transcript may have multiple annotated variants, but only those matching specified filters are retained. The variant sizes listed above for filtering are defaults only and may be adjusted in MINTIE.

### Selecting appropriate control samples

Control samples ideally have the same transcriptional profile as the case samples but without the variants of interest. Controls can be from normal samples from the same tissue type as the case sample if they still have a similar transcriptional program as the case. As RNA sequencing data from the given tissue may be difficult to obtain, other samples from the same cancer type (and same tissue type) can be used as controls. We recommend using at least one control, but ideally 3-10 (using large numbers of controls, >20 for example, will result in increased processing time). In some cases, no appropriate controls may be available, in which case MINTIE can be run without controls. In this mode, differential expression is not performed, however, novel ECs are still selected based on read quantification.

### Simulations

We simulated 1,500 variants using code available in the MINTIE source code repository (https://github.com/Oshlack/MINTIE/tree/master/simu/run_simu.py). Simulations were generated by extracting sequences from the transcripts listed in the hg38 UCSC RefSeq reference, and simulating reads from the resulting sequence. 100 variants from 15 variant types were generated (five fusion types: canonical, extended exon, novel exon, with insertion and unpartnered, five TSV types: insertions, deletions, ITDs, PTDs and inversions, and five novel splice variants: extended exons, novel exons, truncated exons, skipped exons and retained introns. Only transcripts from genes that did not overlap any other genes were used in the simulation. Additionally, each transcript had to have at least 3 exons to be considered as a simulation transcript.

All fusions were simulated by selecting the first two and the last two exons from two random transcripts from different genes, and inserting the intervening sequence. Canonical fusions contained no intervening sequence, while fusions with extended exons inserted 30-199bp of intronic sequence from the end of the second exon of the first transcript. Similarly, fusions with novel exons contained intronic sequence 30-199bp downstream with a size of 30-199bp. Non-canonical fusions with insertions were generated by inserting 7-49bp of randomly-generated sequence between the two fusion transcripts.

Small TSVs were generated by inserting, duplicating or deleting sequence within randomly selected exons from randomly selected transcripts. These small variant types were between 7 and 49 base-pairs and had to reside at least 10bp within the exon. Inversions and partial-tandem duplications were generated by selecting 1-3 random exons within a transcript and either inverting or duplicating their sequence in tandem. Extended and novel exons were generated by adding intronic sequence downstream (directly, or with a 30-199bp gap) of a randomly selected exon. To ensure that novel or extended exons did not overlap exons from other transcripts (or downstream exons of the same transcript), each candidate exon was checked for these potential overlaps (which would otherwise result in obfuscation of the variant, or the wrong variant type being created). Novel junction (skipped exon) variants were created by selecting a random pair of exons and checking whether an existing junction existed between them, creating a transcript with this junction if not. Two randomly-selected neighbouring exons were both truncated at their facing ends (end and start respectively) by 30-199bp. Retained introns included sequence from a randomly selected intron from a given transcript that was >30bp. The presence of correct splicing motifs was not considered for the simulation.

In addition to each variant gene, the sequence of the unaltered wild-type gene was added to the simulated case sample’s reference. An additional 100 unaltered background genes were also added to the case sample. A control sample reference was also generated, which included the unaltered wildtype sequence only for all simulated transcripts. ART-illumina^31^ v2.5.8 was run on the corresponding references with 100bp paired-end reads with a fragment size of 300 and coverage of 50x (transcripts thus have an effective coverage of 100x, given the bi-allelic reference containing variant and wildtype transcripts).

The simulated variant fastq files were run through the MINTIE pipeline with default kmers (29, 49 and 69). Minimum read length was set to 80 (for trimming), minimum assembled transcript length was set to 100, and motif checking was disabled. To ensure that setting a fixed low dispersion value did not adversely over-state the results, we also reran the simulations at four extra dispersion levels: 0.3, 0.5, 0.6 and 0.7 (the original being 0.1). Increasing dispersion had minimal effect on the results until a value between 0.6 and 0.7, after which very few ECs are found to be differentially expressed (Supplementary Figure 1B). This indicated that dispersion is unlikely to play a significant role in the observed results, unless the value is abnormally high (>0.6). All other parameters were left at default. Seven other tools were run on the resulting simulation output: TAP^15^ (commit 8940e45), Barnacle^17^ v1.0.4, SQUID v1.5^16^, JAFFA^24^ v1.07, Arriba^26^ v1.1.0, CICERO^18^ v1.3.0, KisSplice^28^ v2.4.0 and StringTie^12^ v1.3.6. TAP was run using the tap2.py pipeline with kmer lengths of 49 and 79, and `--max_diff_splice` set to 4 to bypass motif checking. Other parameters were kept at default. The transcript reference was generated with PAVfinder’s extract_transcript_sequence.py script, using the UCSC RefGene sequence reference as input. Barnacle was run with its default hg19 reference. SQUID was run using STAR’s alignments. JAFFA was run using its direct mode and default reference. References used in generating simulations were used whenever possible. Default parameters were used for each tool unless otherwise specified. Samtools^59^ v1.8, BLAT^44^ v36.1, STAR^60^ v2.5.3a and bwa^61^ v0.7.17 were used by all methods that required these tools. CICERO was run through the St. Jude Cloud Genomics Platform, using fastq files as input (STAR alignments were performed through St. Jude’s Rapid RNA-seq pipeline, which incorporates CICERO).

Variants were counted as true positives for all tools run for the benchmarking analysis if a variant was reported in the same gene name as the simulated variant (for MINTIE, Arriba and TAP). Gene coordinates were used for all other tools where gene names were not reported (SQUID, StringTie and KissSplice) or where using specific references proved prohibitive (Barnacle). Only transcripts marked as novel were considered for StringTie. Sequence results from KisSplice were aligned with GMAP^58^ (2019-05-12), and aligned coordinates were extracted in order to obtain variant positions. As long as a variant coordinate was found within the expected gene region, this was counted as a hit. Only one gene/coordinate match was needed (of a partnered fusion pair) to be called a true positive. As Barnacle and CICERO were run using a hg19 reference, we used UCSC liftover (https://genome.ucsc.edu/cgi-bin/hgLiftOver) to convert the hg38 simulation coordinates to hg19 (6 TSVs/NSVs and 9 fusion gene coordinates were partially deleted in the liftover; the TSVs/NSVs could not be counted, while the fusion were only affected at one coordinate and could still be counted). All code used to process each tool’s results can be found in the paper analysis code (see Code Availability).

To perform experiments with varying coverage, the simulated fastq files were downsampled using seqtk v1.0 (https://github.com/lh3/seqtk) at 40x, 20x and 10x (resulting in variant coverage of half of each of these values, due to the heterogeneous nature of the simulation). The varying dispersion experiments were generated by manually changing the fixed dispersion parameter (0.1) in the MINTIE source code, to values 0.7, 0.6, 0.5, 0.3. No other parameters were modified. Detection limit experiments were generated by generating nine simulations of 300 variants each (deletions, insertions and ITDs), each with the same random seed (123), only varying by the variant size, ranging from 1 to 9.

### Running MINTIE on Leucegene samples

MINTIE was run on four Leucegene cohorts: the NUP98-NSD1 fusion cohort^62^, the CBF AML cohort^63^, the KMT2A-PTD cohort^32^ and the non-cancer peripheral blood (normals) data set^34^. We used the patient IDs identified by Audemard et al.^64^ to identify the samples containing KMT2A-PTDs, as these were not disclosed in the original paper. For all AML data sets, we used 13 Leucegene normals as the controls (5 monocytes, 5 granulocytes and 3 TWBCs). We also ran the KMT2A-PTD data set against a set of randomly selected AML samples from the CBF cohort (see Supplementary Note 1 for the control samples used). The normal samples were run individually against the remaining samples from that cell type. (The pooled peripheral CD34+ blood cells were not considered for this analysis.) In the granulocyte 1-4 control experiments, all combinations of cases and controls were run for the five total samples, resulting in four total runs for 1 vs. 1, three total runs for 1 vs. 2, two total runs for 1 vs. 3 and a single run for 1 vs. 4.

Salmon^52^ (v0.14) was run separately on the Leucegene normals, the KMT2A-PTD cohort and the selected CBF AML control samples in paired-end mode with default parameters and the --seqBias flag using the same CHESS transcriptome reference used in all other analyses. Counts were summarised at the gene level using tximport^65^ (v1.12.3) with the countsFromAbundance=‘lengthScaledTPM’ parameter. The voom function from Limma^66^ v3.42.0 was used to normalise libraries, and the base prcomp function (R v3.6.2) was used to perform principal component analysis.

Squid and Arriba were run on the Leucegene samples using the same parameters and workflow as on the simulations. As the experiment emphasised sensitivity and compared fewer tools, variants were checked with greater stringency, requiring, in the case of fusions, both variant genes or loci to be called in one variant call (or contig), or as an unknown soft-clip variant containing either fusion gene (in MINTIE’s case). This only happened in two cases, and these variants were manually validated by BLATing the contig sequence to ensure the correct fusion was being reported. In cases where a variant was contained in a single gene and caller output listed gene 1 and gene 2, these both had to match the single gene. A variant filter was also used where classifications were reported (MINTIE, Arriba and TAP) to ensure variants were plausibly classified. Plausible classifications for fusions included ‘fusion’ or ‘unknown’ for MINTIE, ‘translocations’ or ‘inversions’ for Arriba, and ‘fusions’ for TAP. Plausible classifications for ITDs included ‘insertion’, ‘unknown’ or ‘intergenic rearrangement’ for MINTIE, ‘ITD’ or ‘duplication’ for Arriba and ‘ITD’ for TAP. Plausible classifications for PTDs included ‘intergenic rearrangement’ or ‘unknown’ for MINTIE, and ‘duplication’ for Arriba and TAP.

### Running MINTIE on B-ALL and TCGA samples

MINTIE samples used in this study were obtained from the Royal Children’s Hospital; data is described in Brown et al.^35^. Seven samples were excluded due to significantly different sequencing characteristics (shorter read lengths and using an unstranded protocol). Patient samples were selected as controls if they had a positive molecular test and the driver fusions were confirmed in the RNA-seq data. If there were multiple samples obtained from the same patient, they were not considered as controls. This resulted in 34 controls (see Supplementary Note 2 for a list). MINTIE was run with a logFC > 5 cut-off. The RB1 variants were detected in two samples initially, and a third was found after rerunning the pipeline at a lower logFC cutoff (> 2). For the TCGA samples that were identified to contain the variant (using SeqOthello), we obtained the RNA-seq sequence data and ran these samples through the MINTIE pipeline. We used single random TCGA control of the same disease type, sex and similar read length. LogFC cutoff was 2 and kmer sizes of 29 and 39 were used due to short read lengths (samples had 51bp and 76bp reads for the BRCA and ESCA samples respectively).

### Running MINTIE on rare disease samples

MINTIE was run on each of the 52 available samples described in Cummings et al.^14^ against a set of 10 muscle sample controls from GTeX^43^. Due to the relatively short read size (76bp), we selected k-mer sizes of 29, 49 and 69 for SOAPdenovo-Trans. Motif checking was turned off due to the presence of SNP-induced splice sites described in the study. We manually inspected variants called and transcripts assembled in order to determine whether a given variant was called. In order to look for novel candidate variants, we considered all available samples that did not have RNA variants described in the Cummings et al. paper. We applied the following additional filters to the results: i) VAF > 0.1, ii) total reads in controls < 10, iii) < 10 variants found on the de novo assembled transcript, iv) the variant was classified as a TSV or (any type of) fusion, and v) the affected variant affected a gene within the NMD gene list used by the Cummings et al. authors. Each variant was then manually curated considering contig alignment, read coverage and variant location.

## Code availability

- MINTIE software: https://github.com/Oshlack/MINTIE.
- Code for generating paper analyses and figures: https://github.com/Oshlack/MINTIE-paper-analysis (which can be viewed at https://oshlacklab.com/MINTIE-paper-analysis).

## Data availability

- The 1,500 variant simulation (15 variant types), including the downsampled versions, are available under DOI 10.5281/zenodo.4876713, or can be reproduced using the run_simu.py script under https://github.com/Oshlack/MINTIE/tree/master/simu, using the same random seeds as the fullsimu_params.ini file. The downsampled simulations can be reproduced by using seqtk v1.0 with random seeds of 1587, 6471 and 9505 for the 40x, 20x and 10x simulations respectively.
- The 2,700 simulation for testing INDEL and ITD detection limits can be found under DOI 10.5281/zenodo.4876678, and can be reproduced using the code mentioned above (with variant sizes and numbers adjusted as mentioned in the Methods).
- The Leucegene cohorts are available on the Sequence Read Archive: the NUP98-NSD1 fusion cohort^62^ (GSE49642, GSE67039 and GSE52656), the CBF AML cohort^63^ (GSE49642, GSE52656, GSE62190, GSE66917 and GSE67039), the KMT2A-PTD cohort^32^ (accessions GSE49642, GSE52656, GSE66917 and GSE67039) and the non-cancer peripheral blood (normals) data set^34^ (accession GSE51984).
- The paediatric B-ALL samples are available from the European Genome-Phenome Archive (accession number EGAS00001004212).
- The rare disease and control data are available from dbGaP under accession IDs phs000655.v3.p1 and phs000424.v6.p1 (SRR809444, SRR810225, SRR811771, SRR813656, SRR813983, SRR809595, SRR810249, SRR812773, SRR813802 and SRR815020) respectively.
- TCGA samples were obtained from dbGaP under accession ID phs000178
  - Case IDs: 71c5ab4f-ce13-432d-9a90-807ec33cf891 (BRCA) and eae803bf-172f-492f-a381-8f4c040232a2 (ESCA)
  - Control IDs: b5e182ff-159a-44af-881e-8f21bbe96193 (BRCA) and 241a9e1b-3f1a-4eca-90be-d5d48fedce6d (ESCA)

## Abbreviations

SVs: structural variants
TSVs: transcribed structural variants
NSVs: novel splice variants
ECs: equivalence classes
ITD: internal tandem duplication
PTD: partial tandem duplication
WGS: whole genome sequencing
TCGA: The Cancer Genome Atlas
AML: acute myeloid leukaemia
B-ALL: B-cell acute lymphoblastic leukaemia
VAF: variant allele frequency
NTS: non-templated sequence

## Acknowledgements

We would like to thank Jinze Liu and Xiaofei Zhang for querying TCGA with SeqOthello for us, and help from Christoffer Flensburg on interpreting variants and software suggestions. Beryl Cummings, Leigh Waddell, Sandra Cooper and Daniel MacArthur for correspondence about the rare disease data. Some results presented in this manuscript used data generated by the TCGA Research Network: https://www.cancer.gov/tcga. Computing resources were provided by MCRI, The Peter MacCallum Cancer Centre and The University of Melbourne Science IT. This work was funded by NHMRC project grant APP1140626.

## Authors’ contributions

MC: Formal Analysis, Methodology, Software, Visualization, Writing – Original Draft Preparation, Writing – Review & Editing; BS: Visualization; IJM: Writing – Review & Editing, PGE: Writing – Review & Editing; AO: Conceptualization, Supervision, Methodology, Writing – Original Draft Preparation, Writing – Review & Editing; NMD: Conceptualization, Supervision, Methodology, Software, Writing – Original Draft Preparation, Writing – Review & Editing.

## References

1. Saito, M. et al. Development of Lung Adenocarcinomas with Exclusive Dependence on Oncogene Fusions. Cancer Res. 75, 2264–2272 (2015).

2. Patch, A. et al. Whole-genome characterization of chemoresistant ovarian cancer. Nature 489–494 (2015) doi:10.1038/nature14410.

3. Grimwade, D. et al. Refinement of cytogenetic classification in AML Younger adult patients treated in UKMRC. Blood 116, 354–366 (2010).

4. Li, Y. et al. Patterns of somatic structural variation in human cancer genomes. Nature 578, 112–121 (2020).

5. Sanchis-Juan, A. et al. Complex structural variants in Mendelian disorders: identification and breakpoint resolution using short- and long-read genome sequencing. Genome Med. 10, 95 (2018).

6. Holt, J. M. et al. Identification of Pathogenic Structural Variants in Rare Disease Patients through Genome Sequencing. bioRxiv 627661 (2019) doi:10.1101/627661.

7. Cancer, T. et al. Genomic basis for RNA alterations in cancer. Nature 578, 129–136 (2020).

8. Haas, B. J. et al. Accuracy assessment of fusion transcript detection via read-mapping and de novo fusion transcript assembly-based methods. Genome Biol. 20, 1–16 (2019).

9. Kumar, A. et al. Substantial interindividual and limited intraindividual genomic diversity among tumors from men with metastatic prostate cancer. Nat. Med. 22, 1–13 (2016).

10. Trapnell, C. et al. Differential gene and transcript expression analysis of RNA-seq experiments with TopHat and Cufflinks. Nat. Protoc. 7, 562–78 (2012).

11. Sacomoto, G. A. T. et al. KISSPLICE: de-novo calling alternative splicing events from RNA-seq data. BMC Bioinformatics 13, 1–12 (2012).

12. Pertea, M. et al. StringTie enables improved reconstruction of a transcriptome from RNA-seq reads. Nat. Biotechnol. 33, 290–295 (2015).

13. Gonorazky, H. D. et al. Expanding the Boundaries of RNA Sequencing as a Diagnostic Tool for Rare Mendelian Disease. Am. J. Hum. Genet. 104, 1007 (2019).

14. Cummings, B. B. et al. Improving genetic diagnosis in Mendelian disease with transcriptome sequencing. Sci. Transl. Med. 9, (2017).

15. Chiu, R., Nip, K. M., Chu, J. & Birol, I. TAP: a targeted clinical genomics pipeline for detecting transcript variants using RNA-seq data. BMC Med. Genomics 11, 79 (2018).

16. Ma, C., Shao, M. & Kingsford, C. SQUID: Transcriptomic structural variation detection from RNA-seq. Genome Biol. 19, 1–16 (2018).

17. Swanson, L. et al. Barnacle: detecting and characterizing tandem duplications and fusions in transcriptome assemblies. BMC Genomics 14, 550 (2013).

18. Tian, L. et al. CICERO: a versatile method for detecting complex and diverse driver fusions using cancer RNA sequencing data. Genome Biol. 21, 126 (2020).

19. Mullighan, C. G. et al. Deletion of IKZF1 and Prognosis in Acute Lymphoblastic Leukemia. N. Engl. J. Med. 360, 470–480 (2009).

20. Bolouri, H. et al. The molecular landscape of pediatric acute myeloid leukemia reveals recurrent structural alterations and age-specific mutational interactions. Nat. Med. (2017) doi:10.1101/125609.

21. An integrated map of structural variation in 2,504 human genomes | Nature. https://www.nature.com/articles/nature15394.

22. STAR-Fusion: Fast and Accurate Fusion Transcript Detection from RNA-Seq | bioRxiv. https://www.biorxiv.org/content/10.1101/120295v1.abstract.

23. Kim, D. & Salzberg, S. L. TopHat-Fusion: An algorithm for discovery of novel fusion transcripts. Genome Biol. 12, (2011).

24. Davidson, N. M., Majewski, I. J. & Oshlack, A. JAFFA: High sensitivity transcriptome-focused fusion gene detection. Genome Med. 7, 43 (2015).

25. Melsted, P. et al. Fusion detection and quantification by pseudoalignment. bioRxiv 166322 (2017) doi:10.1101/166322.

26. Uhrig, S. et al. Accurate and efficient detection of gene fusions from RNA sequencing data. Genome Res. gr.257246.119 (2021) doi:10.1101/gr.257246.119.

27. Qiu, Y., Ma, C., Xie, H. & Kingsford, C. Detecting transcriptomic structural variants in heterogeneous contexts via the Multiple Compatible Arrangements Problem. Algorithms Mol. Biol. 15, 9 (2020).

28. Sacomoto, G. A. T. et al. KISSPLICE: de-novo calling alternative splicing events from RNA-seq data. BMC Bioinformatics 13, 1–12 (2012).

29. Audoux, J. et al. DE-kupl: exhaustive capture of biological variation in RNA-seq data through k-mer decomposition. Genome Biol. 18, 243 (2017).

30. O’Leary, N. A. et al. Reference sequence (RefSeq) database at NCBI: current status, taxonomic expansion, and functional annotation. Nucleic Acids Res. 44, D733–745 (2016).

31. Huang, W., Li, L., Myers, J. R. & Marth, G. T. ART: a next-generation sequencing read simulator. Bioinformatics 28, 593–594 (2012).

32. Lavallée, V.-P. et al. The transcriptomic landscape and directed chemical interrogation of MLL-rearranged acute myeloid leukemias. Nat. Genet. 47, 1030–1037 (2015).

33. Audemard, É. et al. Target variant detection in leukemia using unaligned RNA-Seq reads. bioRxiv 295808 (2018) doi:10.1101/295808.

34. Pabst, C. et al. GPR56 identifies primary human acute myeloid leukemia cells with high repopulating potential in vivo. Blood 127, 2018–2027 (2016).

35. Brown, L. M. et al. The application of RNA sequencing for the diagnosis and genomic classification of pediatric acute lymphoblastic leukemia. 4, 1–3 (2020).

36. Gröbner, S. N. et al. The landscape of genomic alterations across childhood cancers. Nature 555, 321–327 (2018).

37. Ma, X. et al. Pan-cancer genome and transcriptome analyses of 1,699 paediatric leukaemias and solid tumours. Nat. Publ. Group (2018) doi:10.1038/nature25795.

38. Mullighan, C. G. et al. Deletion of IKZF1 and Prognosis in Acute Lymphoblastic Leukemia. N. Engl. J. Med. 360, 470–480 (2009).

39. Mullighan, C. G. et al. Genome-wide analysis of genetic alterations in acute lymphoblastic leukaemia. Nature 446, 758–764 (2007).

40. Gu, Z. et al. PAX5-driven subtypes of B-progenitor acute lymphoblastic leukemia. Nat. Genet. doi:10.1038/s41588-018-0315-5.

41. Zhang, J. et al. Key pathways are frequently mutated in high-risk childhood acute lymphoblastic leukemia: a report from the Children’s Oncology Group. Blood 118, 3080–3087 (2011).

42. Yu, Y. et al. SeqOthello: Query over RNA-seq experiments at scale. bioRxiv 258772 (2018) doi:10.1101/258772.

43. Consortium, T. Gte. The Genotype-Tissue Expression (GTEx) pilot analysis: Multitissue gene regulation in humans. Science 348, 648–660 (2015).

44. BLAT—The BLAST-Like Alignment Tool. https://genome.cshlp.org/content/12/4/656.abstract.

45. Waddell, L. B. et al. WGS and RNA Studies Diagnose Noncoding DMD Variants in Males With High Creatine Kinase. Neurol. Genet. 7, e554 (2021).

46. Xu, H. et al. FastUniq: A Fast De Novo Duplicates Removal Tool for Paired Short Reads. PLOS ONE 7, e52249 (2012).

47. Bolger, A. M., Lohse, M. & Usadel, B. Trimmomatic: a flexible trimmer for Illumina sequence data. Bioinformatics 30, 2114–2120 (2014).

48. Xie, Y. et al. SOAPdenovo-Trans: De novo transcriptome assembly with short RNA-Seq reads. Bioinformatics 30, 1660–1666 (2014).

49. Haas, B. J. et al. De novo transcript sequence reconstruction from RNA-Seq: reference generation and analysis with Trinity. Nat Protocols vol. 8 (2014).

50. Bushmanova, E., Antipov, D., Lapidus, A. & Prjibelski, A. D. rnaSPAdes: a de novo transcriptome assembler and its application to RNA-Seq data. GigaScience 8, giz100 (2019).

51. Pertea, M. et al. CHESS: a new human gene catalog curated from thousands of large-scale RNA sequencing experiments reveals extensive transcriptional noise. Genome Biol. 19, 332825 (2018).

52. Patro, R., Duggal, G., Love, M. I., Irizarry, R. A. & Kingsford, C. Salmon provides fast and bias-aware quantification of transcript expression. Nat. Methods 14, 021592 (2017).

53. Robinson, M. D., McCarthy, D. J. & Smyth, G. K. edgeR: a Bioconductor package for differential expression analysis of digital gene expression data. Bioinforma. Oxf. Engl. 26, 139–40 (2010).

54. Chen, Y., Lun, A. T. L. & Smyth, G. K. Differential Expression Analysis of Complex RNA-seq Experiments Using edgeR. in Statistical Analysis of Next Generation Sequencing Data (eds. Datta, S. & Nettleton, D.) 51–74 (Springer International Publishing, 2014). doi:10.1007/978-3-319-07212-8_3.

55. Lund, S. P., Nettleton, D., McCarthy, D. J. & Smyth, G. K. Detecting differential expression in RNA-sequence data using quasi-likelihood with shrunken dispersion estimates. Stat. Appl. Genet. Mol. Biol. 11, (2012).

56. Phipson, B., Lee, S., Majewski, I. J., Alexander, W. S. & Smyth, G. K. ROBUST HYPERPARAMETER ESTIMATION PROTECTS AGAINST HYPERVARIABLE GENES AND IMPROVES POWER TO DETECT DIFFERENTIAL EXPRESSION. Ann. Appl. Stat. 10, 946–963 (2016).

57. McCarthy, D. J., Chen, Y. & Smyth, G. K. Differential expression analysis of multifactor RNA-Seq experiments with respect to biological variation. Nucleic Acids Res. 40, 4288–4297 (2012).

58. Wu, T. D., Reeder, J., Lawrence, M., Becker, G. & Brauer, M. J. GMAP and GSNAP for Genomic Sequence Alignment: Enhancements to Speed, Accuracy, and Functionality. in Statistical Genomics: Methods and Protocols (eds. Mathé, E. & Davis, S.) 283–334 (Springer, 2016). doi:10.1007/978-1-4939-3578-9_15.

59. Li, H. et al. The Sequence Alignment/Map format and SAMtools. Bioinforma. Oxf. Engl. 25, 2078–9 (2009).

60. Dobin, A. et al. STAR: Ultrafast universal RNA-seq aligner. Bioinformatics 29, 15–21 (2013).

61. Li, H. & Durbin, R. Fast and accurate short read alignment with Burrows-Wheeler transform. Bioinforma. Oxf. Engl. 25, 1754–1760 (2009).

62. Lavallée, V. P. et al. Identification of MYC mutations in acute myeloid leukemias with NUP98-NSD1 translocations. Leukemia 30, 1621–1624 (2016).

63. Thompson, M. P., Waters, T. M., Kaplan, E. K., Mckillop, C. N. & Martin, M. G. RNA-sequencing analysis of core binding factor AML identifies recurrent ZBTB7A mutations and defines RUNX1-CBFA2T3 fusion signature. 128, 872–875 (2016).

64. Audemard, É. et al. Target variant detection in leukemia using unaligned RNA-Seq reads. bioRxiv 295808 (2018) doi:10.1101/295808.

65. Soneson, C., Love, M. I. & Robinson, M. D. Differential analyses for RNA-seq: transcript-level estimates improve gene-level inferences. F1000Research 4, 1521 (2016).

66. Ritchie, M. E. et al. limma powers differential expression analyses for RNA-sequencing and microarray studies. Nucleic Acids Res. 43, e47–e47 (2015).

