## Supplementary Material for "MINTIE: identifying novel structural and splice variants in transcriptomes using RNA-seq data"

### Supplementary Figures

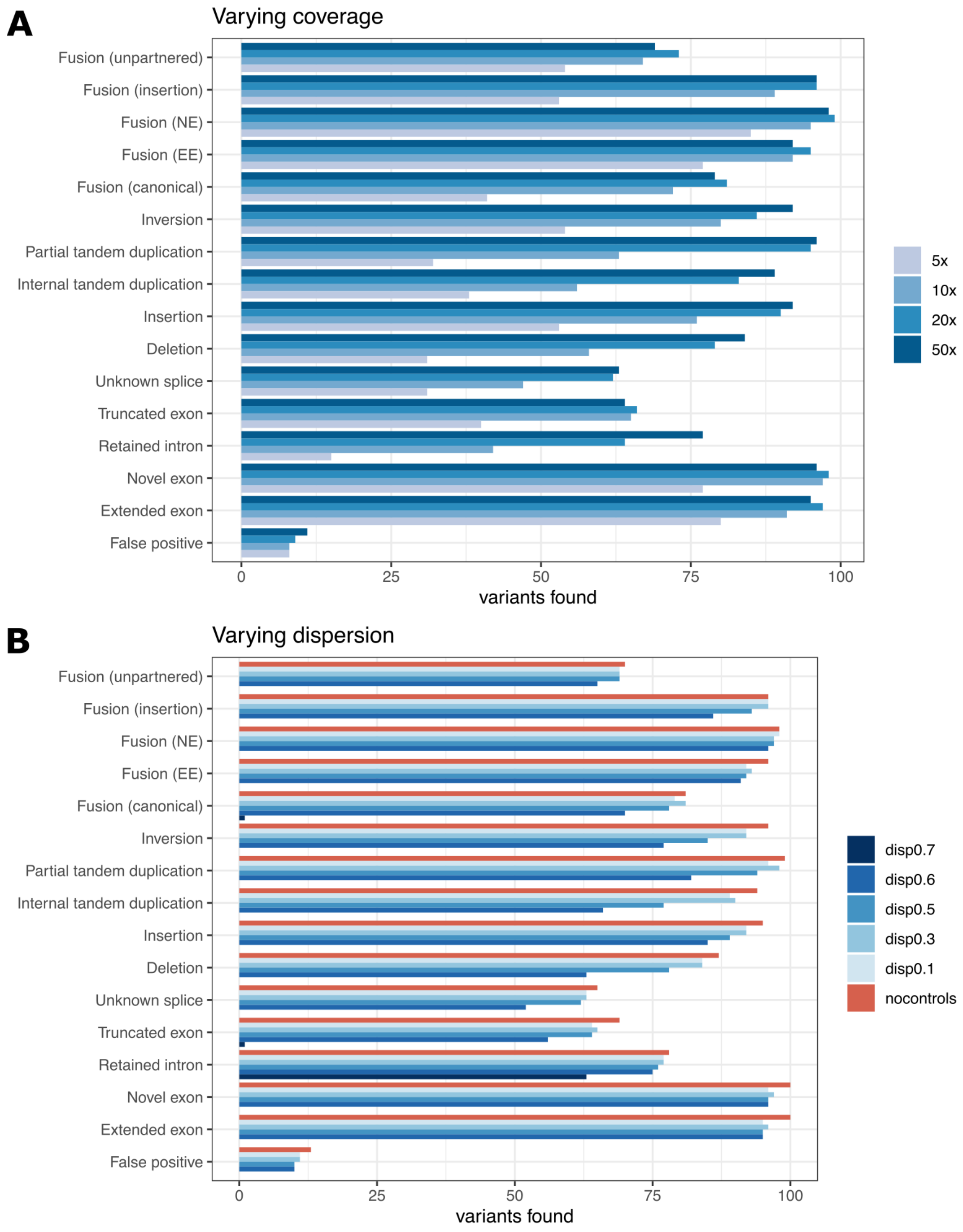


***Supplementary Figure 1** | Variants detected in simulations with varying coverage **(A)** and varying dispersion **(B)**.*

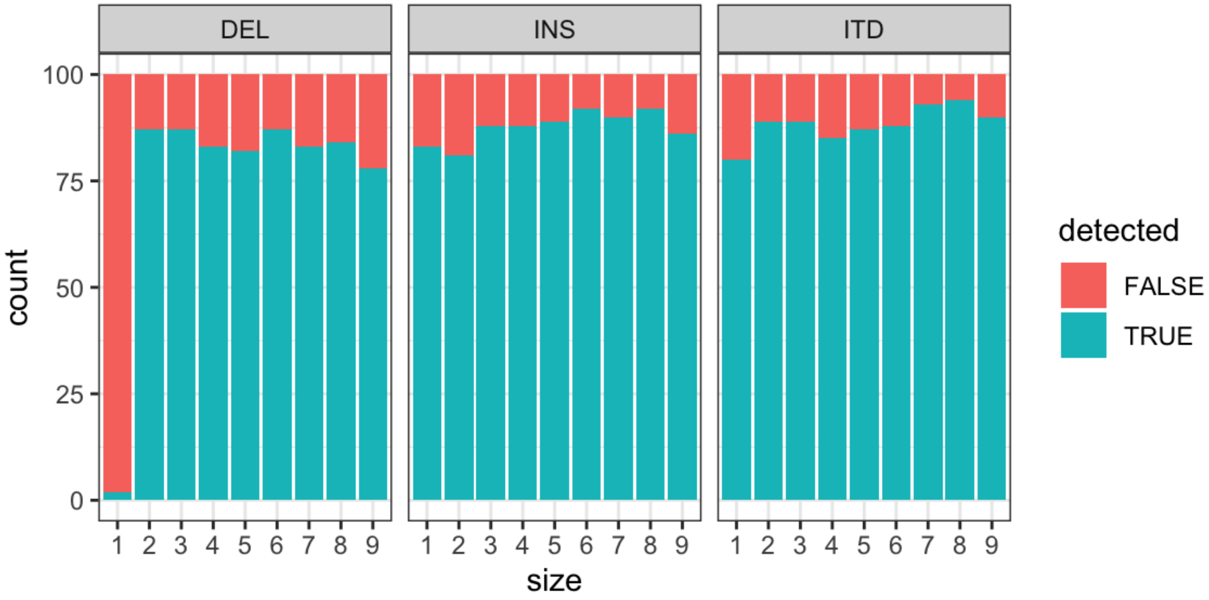


***Supplementary Figure 2*** | Number of deletions, insertions and ITDs detected by MINTIE at variant sizes 1-9, with 100 variants generated per variant type and size category.

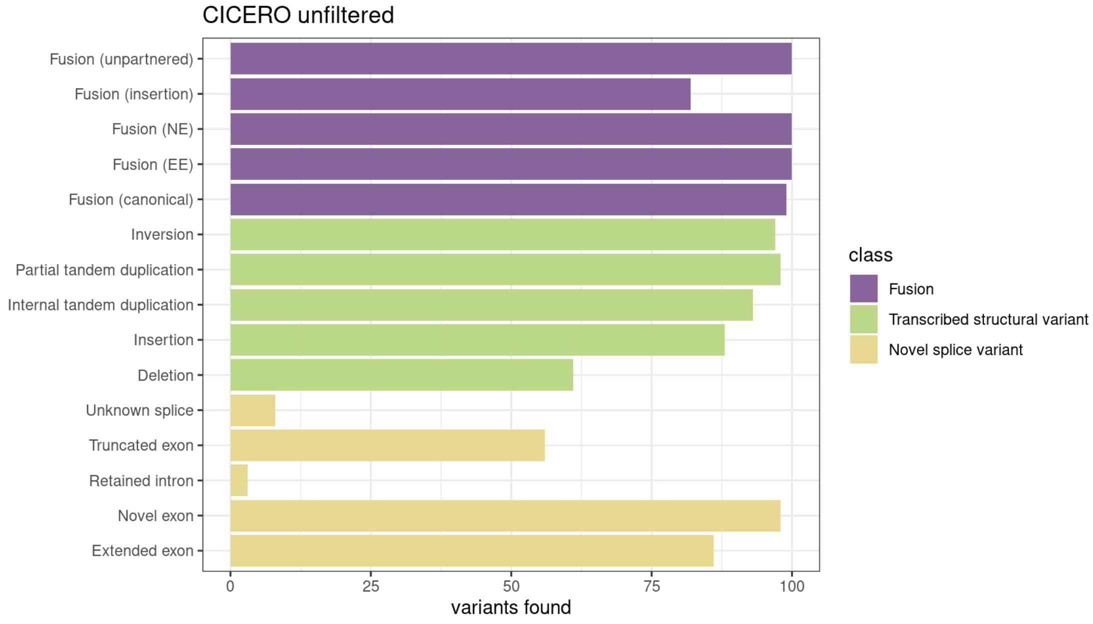


***Supplementary Figure 3*** | *Results of the unfiltered output from the CICERO fusion caller on the simulated data. In addition, 596 false positive calls were identified in the unfiltered output (not shown).*

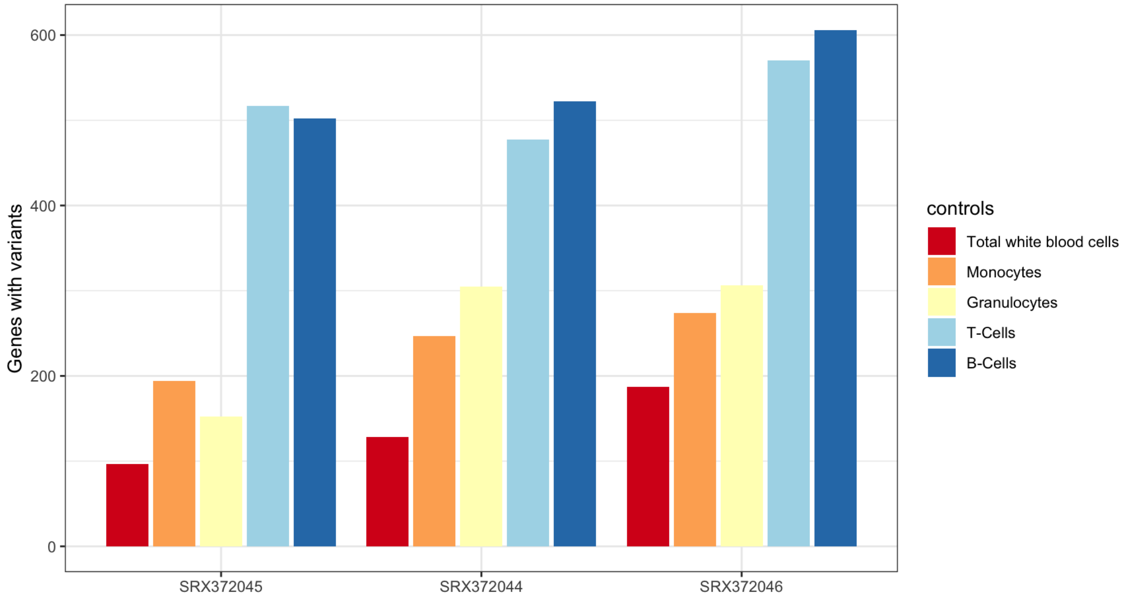


***Supplementary Figure 4 |*** *Number of genes with variants found in three Leucegene total white blood cell (TWBC) samples with different normal cell types as controls. Controls are ordered by total variants found across all three samples with controls of the same type (TWBCs) resulting in the fewest variant genes. TWBCs consisted of two control samples, with all other control types consisting of five.*

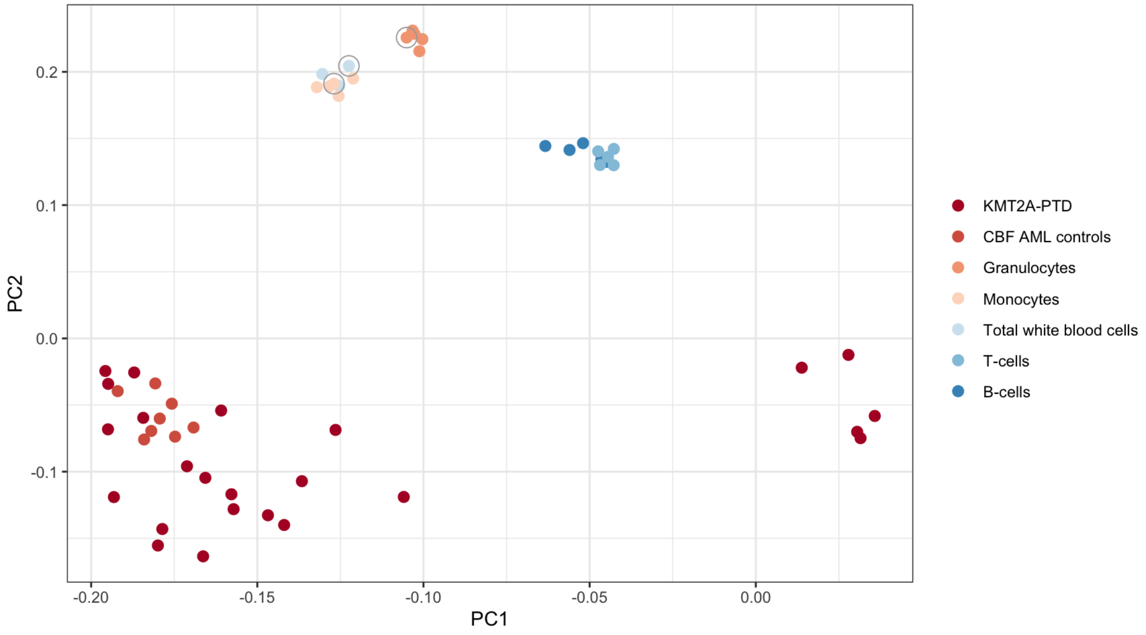


***Supplementary Figure 5 |*** *PCA plot of KMT2A-PTD cohort, compared with selected CBF AML controls, and Leucegene normals (granulocytes, monocytes, total white blood cells, T-cells and B-cells), derived from Salmon quantification of the top 500 most variable genes. The reduced control set is circled.*

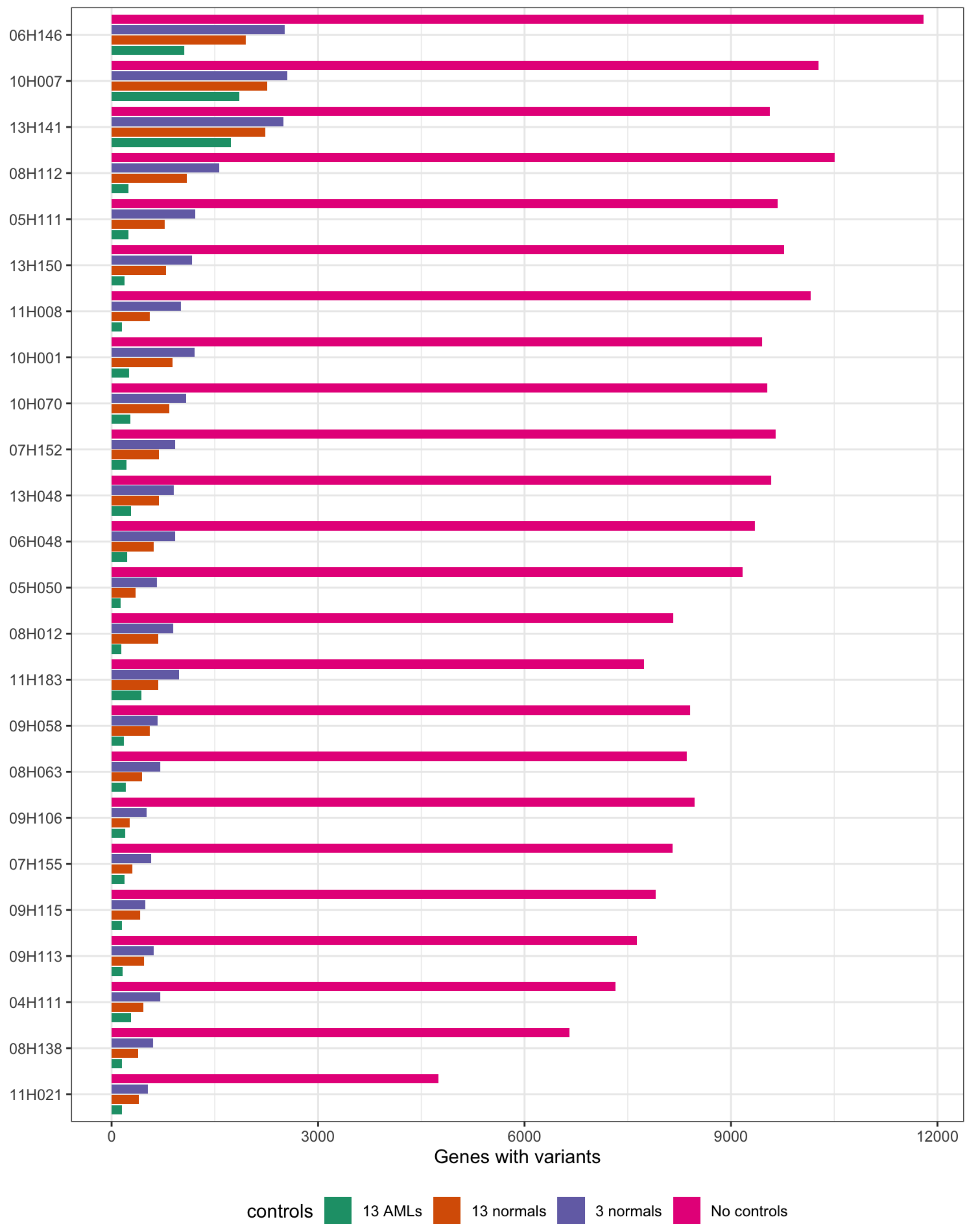


**Supplementary Figure 6 |** *Number of variant genes found in the KMT2A-PTD cohort (24 Leucegene samples containing KMT2A alterations) when using three different Leucegene control groups: a randomly selected set of 13 AMLs, 13 normals, 3 normals (subset of the 13) and no controls.*

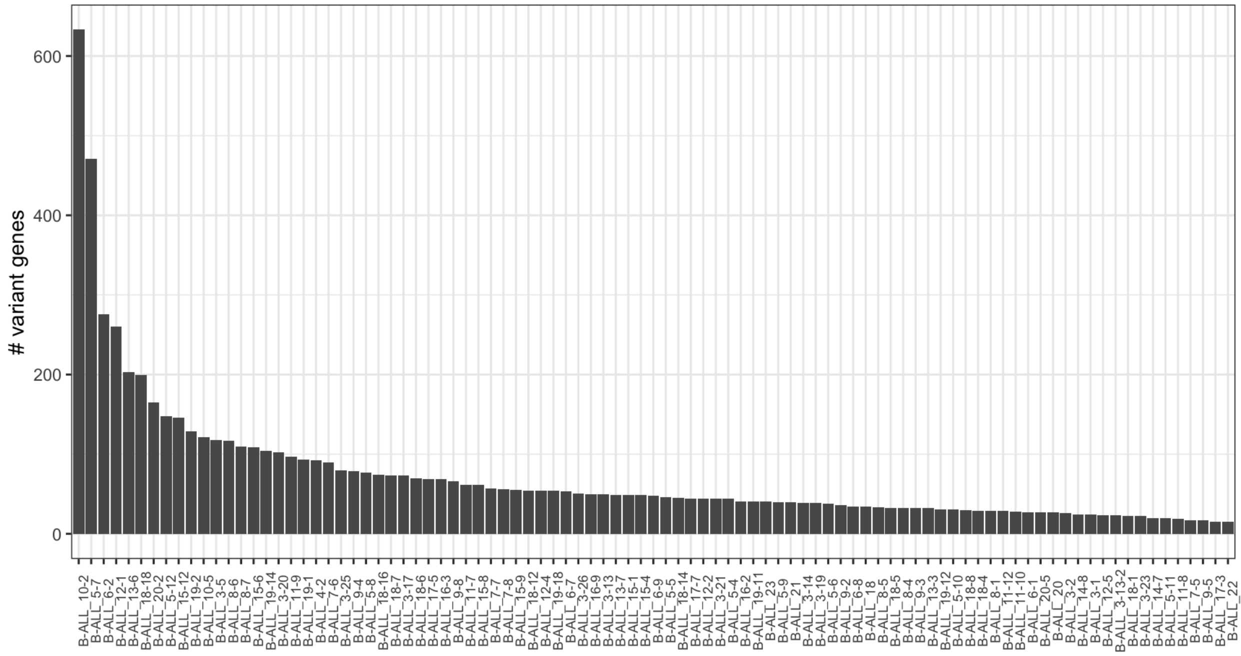


***Supplementary Figure 7 |*** *Number of variant genes found per sample in RCH B-ALL cohort.*

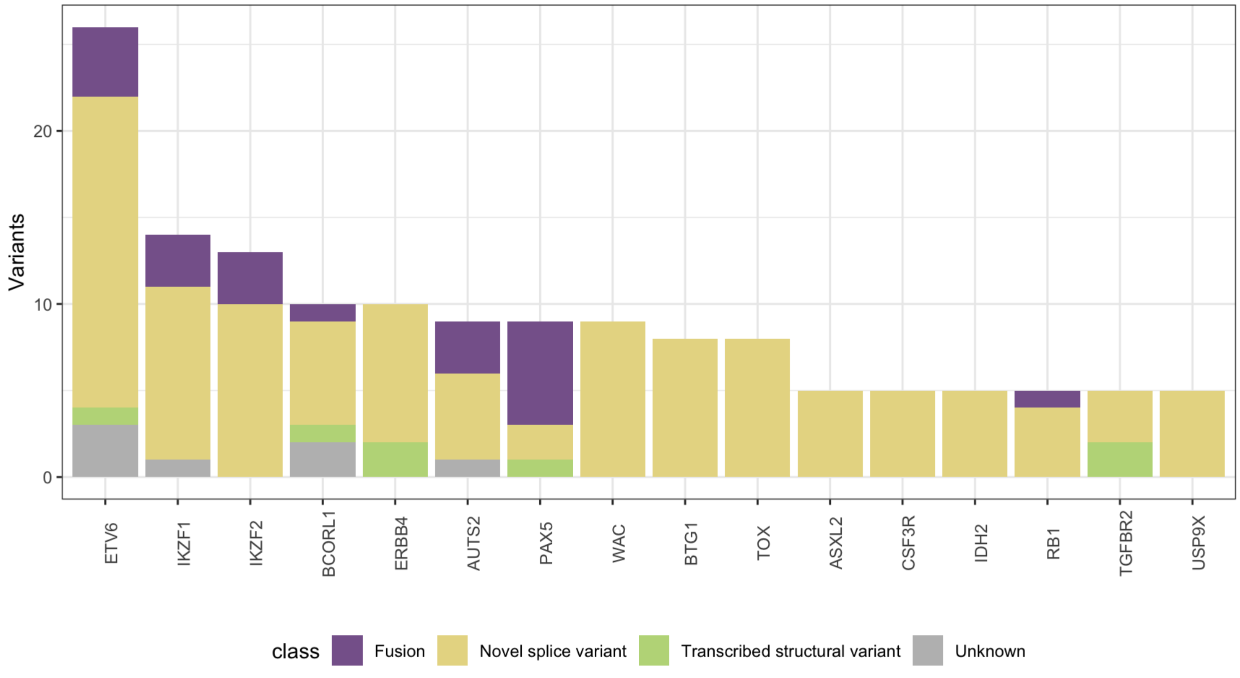


***Supplementary Figure 8 |*** *The number of variants found in ALL-associated genes across the RCH B-ALL cohort.*

### Supplementary Tables

***Supplementary Table 1 |*** *Validated KMT2A-PTDs from a prior study detected by MINTIE in a 24-sample Leucegene cohort run against 3 Leucegene normals (one each of monocytes, granulocytes and total white blood cells), 13 Leucegene normals (5 monocytes, 5 granulocytes and 3 total white blood cells), a randomly selected cohort of 13 Leucegene CBF AMLs with no known KMT2A rearrangement and no controls. Coverage obtained from Audemard et al. In cases where multiple PTDs were detected in the same patient, the highest min coverage was used.*

| Patient | 3 normals | 13 normals | 13 AMLs | No controls | Coverage |
|---|---|---|---|---|---|
| 07H152 | Y | Y | N | Y | 158 |
| 09H115 | Y | Y | N | Y | 125 |
| 06H146 | Y | Y | Y | Y | 87 |
| 05H111 | Y | Y | Y | Y | 79 |
| 11H021 | Y | Y | N | Y | 63 |
| 05H050 | Y | Y | Y | Y | 58 |
| 13H150 | N | Y | N | Y | 58 |
| 13H048 | Y | Y | Y | Y | 57 |
| 10H070 | Y | Y | Y | Y | 53 |
| 10H007 | Y | Y | Y | Y | 50 |
| 09H106 | Y | Y | Y | Y | 49 |
| 13H141 | Y | Y | Y | Y | 45 |
| 09H058 | Y | Y | Y | Y | 29 |
| 07H155 | Y | Y | N | Y | 23 |
| 08H112 | Y | Y | Y | Y | 22 |
| 09H113 | Y | Y | Y | Y | 17 |
| 11H183 | N | N | N | N | 16 |
| 08H012 | N | N | N | N | 15 |
| 08H138 | N | N | N | Y | 15 |
| 11H008 | N | N | N | Y | 13 |
| 06H048 | N | N | N | Y | 10 |
| 08H063 | N | N | N | N | 6 |
| 10H001 | N | N | N | Y | 6 |
| 04H111 | N | N | N | N | 3 |
| **Found (of 24)** | **11** | **16** | **15** | **19** | |

***Supplementary Table 2*** **|** *Novel variants found in the B-ALL cohort in clinically relevant genes. All locations are hg38. RB1 and ETV6 variants were both detected in 3 samples total.*

| **Gene** | **Variant type** | **Samples affected** | **Novel splice junction** | **Sequence queried by SeqOthello** | **TCGA hit** |
|---|---|---|---|---|---|
| RB1 | Unpartnered fusion | 3 | chr13:48381444-48644164 | TAGAACGATGTGAACATCGAATCATGGAATCCCTTGCATGGCTCTCAAGTCAGTTCCTGCCCCACTGCCCCACAGAAGTGTTTTCTGATGTGCT | |
| RB1 | Unpartnered fusion | 1 | chr13:48381444-48647695 | TAGAACGATGTGAACATCGAATCATGGAATCCCTTGCATGGCTCTCAAGCCTTGTATACACTCAATATGCAAGAAGCCCTGGAAGTTCCCAAGGT | |
| RB1 | Unpartnered fusion | 1 | chr13:48381444-48530084 | TAGAACGATGTGAACATCGAATCATGGAATCCCTTGCATGGCTCTCAAGACAGAGTTTTGCCATGTTGTCCCGGCTGGTCTCTAACTCCTGGGCT | 1 BRCA, 1 ESAD |
| IKZF1 | PTD | 1 | chr7:50382708-50382540 | CTGCCGCCGGAGGGACGCCCTCACTGGCCACCTGAGGACGCACTCCGGAGAACGGCCCTTCCAGTGCAATCAGTGCGGGGCCTCATTCACCCAGA | |
| PAX5 | PTD | 1 | chr9:37020805-37002647 | ATCACGTCCCCCAGCGCCGACACCAACAAGCGCAAGAGAGACGAAGGACATGGAGGAGTGAATCAGCTTGGGGGGGTTTTTGTGAATGGACGGCC | |
| ETV6 | Skipped exons | 1 | chr12:11752580-11869424 | AGCGCTCAGGATGGAGGAAGACTCGATCCGCCTGCCTGCGCACCTGCATAACTGTGTCCAGAGGACCCCCAGGCCATCCGTGGATAATGTGCACC | 1 LSCC |
| ETV6 | Skipped exons | 2 | chr12:11752580-11890941 | AGCGCTCAGGATGGAGGAAGACTCGATCCGCCTGCCTGCGCACCTGCGTTTATGAAAACCCCAGATGAAATCATGAGTGGCCGAACAGACCGTCT | |
| ETV6 | Skipped exons + novel exons | 1 | chr12:11869970-11931173 | TGTCTCCCCGCCTGAAGAGCACGCCATGCCCATTGGGAGAATAGCAGGTCCCATCCCATCCGAGTCTCAACAGAAACATCACCTCCCCAGGGAGA | |

***Supplementary Table 3*** **|** *Novel candidate variants found in the rare disease data.*

| Patient | Genetic Diagnosis | Gene | Location 1 | Location 2 | Size | Variant Type | rs ID |
|---|---|---|---|---|---|---|---|
| C5 | Strong candidate gene | DMD | chrX:32287529 | chr8:65268274, chr8:65267608, chr8:65194163 | N/A | Unpartnered Fusion | |
| N24 | No strong candidates | DMD | chrX:31121428 | chrX:31121443 | 17 | 3' UTR Deletion | rs763028610 |
| N9 | No strong candidates | KLHL9 | chr9:21335327 | chr9:21335339 | 12 | 5' UTR Deletion | rs201092918 |
| N23 | No strong candidates | KLHL9 | chr9:21335327 | chr9:21335339 | 12 | 5' UTR Deletion | rs201092918 |
| D9 | Diagnosed | LDB3 | chr10:86735779 | chr10:86735786 | 7 | 3' UTR Deletion | rs746342719 |
| N24 | No strong candidates | LDB3 | chr10:86735779 | chr10:86735786 | 7 | 3' UTR Deletion | rs746342719 |
| N11 | No strong candidates | LDB3 | chr10:86735779 | chr10:86735786 | 7 | 3' UTR Deletion | rs746342719 |
| D13 | Diagnosed | LDB3 | chr10:86735779 | chr10:86735786 | 7 | 3' UTR Deletion | rs746342719 |
| N6 | No strong candidates | VAPB | chr20:58445685 | chr20:58445701 | 16 | 3' UTR Deletion | rs138225455 |

### Supplementary Notes

***Supplementary Note 1 |*** *List of Leucegene samples analysed.*

Core binding factor cohort: 03H065, 03H083, 03H095*, 03H109, 03H112, 04H030*, 04H061*, 04H091*, 05H042*, 05H099*, 05H113, 05H118*, 05H136*, 05H184*, 06H020, 06H035*, 06H115*, 07H099*, 07H137*, 07H144, 08H034, 08H042, 08H072, 08H081, 08H099, 09H016, 09H040, 09H066, 10H008, 10H030, 10H119, 11H022, 11H104, 11H107, 11H179, 12H042, 12H044, 12H045, 12H098, 12H165, 12H166, 12H180, 12H183, 13H066, 13H120, 13H169 (*used as controls).

NUP98-NSD1 cohort: 03H041, 05H034, 05H163, 08H049, 10H038, 11H027, 11H160.

KMT2A-PTD cohort: 05H050, 09H113, 09H115, 11H021, 08H012, 08H112, 11H008, 05H111, 06H146, 04H111, 06H048, 07H152, 07H155, 08H063, 08H138, 09H058, 09H106, 10H001, 10H007, 10H070, 11H183, 13H048, 13H141, 13H150

***Supplementary Note 2 |*** *Samples used as controls in the RCH B-ALL analysis.*

B-ALL3-1, B-ALL3-2, B-ALL5-8, B-ALL5-10, B-ALL5-11, B-ALL7-7, B-ALL8-1, B-ALL9-2, B-ALL9-5, B-ALL11-8, B-ALL12-4, B-ALL14-7, B-ALL14-8, B-ALL16-2, B-ALL16-3, B-ALL17-3, B-ALL18-14, B-ALL9-8, B-ALL10-2, B-ALL6-9, B-ALL11-10, B-ALL15-9, B-ALL19-11, B-ALL6-1, B-ALL18-4, B-ALL13-7, B-ALL19-12, B-ALL6-8, B-ALL3-26, B-ALL-22, B-ALL16-9, B-ALL3-20, B-ALL3-25, B-ALL18-7
